# Preoptic Neurons that Control Entry into Hibernation

**DOI:** 10.64898/2025.12.05.692394

**Authors:** Adrian J. Martinez, Christopher M. Reid, Aurora J. Lavin-Peter, Wenhui Li, Andrew S. Lee, Eric C. Griffith, Sinisa Hrvatin

**Affiliations:** Whitehead Institute for Biomedical Research, Cambridge, MA 02142, USA; Department of Brain and Cognitive Sciences, Massachusetts Institute of Technology, Cambridge, MA 02142, USA; Department of Neurobiology, Harvard Medical School, Boston, MA 02115, USA; Department of Biology, Massachusetts Institute of Technology, Cambridge, MA 02142, USA; Howard Hughes Medical Institute, Massachusetts Institute of Technology, Cambridge, MA, USA

## Abstract

Evolution of seasonal hibernation has enabled mammals to survive harsh conditions by entering a state of prolonged hypometabolism and hypothermia with body temperatures as low as 0-4°C^1–6^. Despite decades of physiological studies, the genetic tools to study hibernation have remained limited and the mechanisms that induce hibernation entry are still unknown. Focusing on the brain, we map state-dependent neuronal activity across the hibernation cycle in Syrian hamsters and identify the hypothalamic anterior preoptic area (aPOA) as a key regulator of hibernation entry. Single-nucleus RNA and chromatin profiling provided a map of neuronal populations present in the hamster POA and enabled the discovery and design of an enhancer AAV that selectively targets hibernation-associated aPOA subpopulations. Using this genetic approach, we show that *Samd3*-positive aPOA glutamatergic neurons are necessary for entry into hibernation and that their activation is sufficient to induce a prolonged hypothermic state with associated nesting behavior. Together, we identify the first neuronal population that controls entry into hibernation, opening new avenues for investigating and manipulating the metabolic and physiological mechanisms underlying this extreme state of “suspended animation” and its potential applications in aging and disease.

## Main Text

Maintaining core body temperature (T_b_) within a narrow range is a defining feature of warm-blooded animals^7,8^. Yet, to survive in extreme environments, many birds and mammals, including some primates, have evolved the ability to initiate hibernation, a state associated with profound reductions in heart rate, respiration, metabolic rate, and core body temperature—sometimes reaching as low as 0-4°C^1–6^. How warm-blooded animals initiate such extreme hypometabolic states and how their cells survive at these low temperatures have long remained fundamental questions in homeotherm biology. Moreover, an understanding of the mechanisms controlling hibernation could have important medical applications from therapeutic hypothermia and organ preservation to prevention of muscle atrophy, aging, and potentially even space travel^9–20^.

The defining feature of hibernation is a controlled reduction of metabolic rate, yet hibernation is regulated and expressed in distinct ways across species^2,3^. Large hibernators, such as bears, maintain a relatively constant T_b_ near 30-36°C^21^, whereas hibernating rodents like the arctic ground squirrels decrease T_b_ to below 0°C^6^. Animals in moderate and cold climates typically hibernate during winter, while tropical hibernators, such as lemurs, hibernate in the dry warm season^3,22^. Given the phenotypic differences and breadth of animals capable of hibernation, it remains unclear to what extent the underlying mechanisms regulating hibernation are conserved across species.

The search for the triggers of hibernation entry is long-standing, with multiple models having been proposed, including the existence of circulating hibernation-inducing factors^23–26^. More recently, advances in neuroscience have enabled detailed mapping of the core brain–body thermoregulatory circuits^7,8,27–38^ and have identified hypothalamic neuronal populations in mice that regulate fasting-induced daily torpor^39–44^. However, these studies were conducted in non-hibernating species and the extent to which homologous circuits evolved to regulate seasonal hibernation has yet to be determined. Moreover, the lack of genetic access in hibernators has thus far prevented the identification, dissection, and manipulation of brain circuits that regulate seasonal hibernation. As a result, although multiple environmental cues and hormonal systems are known to modulate hibernation timing and depth in different species^1,45–57^, which neuronal populations control entry into hibernation has remained unknown.

To probe the neural mechanisms that control entry into hibernation, we established a laboratory paradigm of hibernation in the Syrian hamster (*Mesocricetus auratus*). In these animals, hibernation is defined by repeated bouts of profoundly decreased metabolic rate and body temperature called *deep torpor* (DT) punctuated by brief arousals that restore normal body temperature referred to as interbout euthermia (IBE)^2,6,58–61^. To monitor entry into DT, we implanted each animal with telemetric temperature probes. To induce hibernation, adult (5-6-week-old) animals housed at room temperature (∼22°C) with standard photoperiod (12 hours light (L): 12 hours dark (D)) were first shifted to a short photoperiod (8L:16D) for 8-10 weeks, followed by transfer to 4-5°C ambient temperature conditions to simulate seasonal changes (Fig. 1a, b, Extended Data Fig. 1a). While this paradigm induced hibernation in both males and females, female hamsters were initially found to hibernate more readily (data not shown), with ∼80% initiating torpor bouts within six weeks of ambient temperature reduction. As a result, we restricted our subsequent analyses to females.

**Fig. 1:**
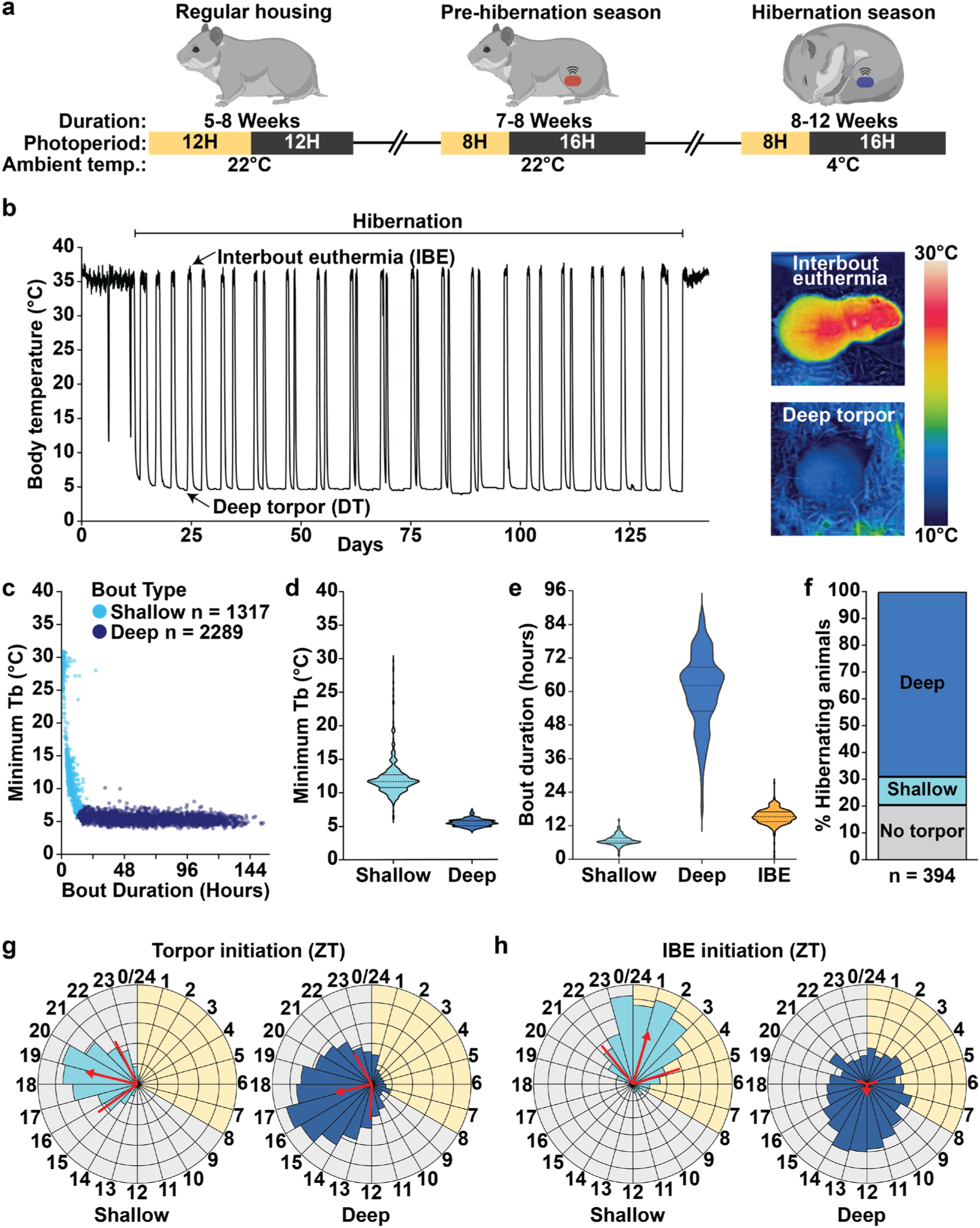
A laboratory model of Syrian hamster hibernation. **a**, A reproducible laboratory model that induces hibernation with stereotyped deep bouts of torpor in Syrian hamsters. **b**, Left: Representative body temperature trace of a hibernating hamster, consisting of repeated bouts of deep torpor (DT) and interbout euthermia (IBE). Right: Thermal images of a hibernating hamster during euthermic and deep torpor intervals. **c**, Classification of torpor bouts as shallow or deep based on minimum temperature and bout duration. **d,e**, Violin plots of minimum body temperature (**d**), and duration (**e**) of shallow and deep torpor bouts as well as IBE periods (n = 2471). **f**, Percentage of animals undergoing DT, only shallow torpor, or no torpor bouts within 6 weeks of initiating the hibernation period. **g**, Circadian representation (Raleigh plot) of the time of entry into shallow (light blue) or deep (dark blue) torpor bouts. Light and dark cycle phases are represented by yellow and grey, respectively. Red arrow direction and length indicate mean time of initiation and concentration around the mean, respectively. Red lines flanking the arrow indicate s.d. **h**, Circadian representation as in (**g**) of entry into IBE following deep (left, n = 1496) or shallow (right, n = 1011) torpor bouts.

While typical torpor-associated T_b_ decreases followed an exponential decay curve (n = 1262 torpor bouts, Extended Data Fig. 1b), animals exhibited some variability in torpor dynamics. Altogether, 69% of animals (n = 271) achieved DT (minimal T_b_ = 5.5 ± 0.5°C, bout duration 60.3 ± 13.4 h) and 11% (n = 42) exhibited only shallow torpor bouts (minimal T_b_ = 12.2 ± 3.1°C, bout duration 6.8 ± 1.8 h), with 20% (n = 81) failing to initiate torpor during the 6-week observation period (Fig. 1c-f, Extended Data Fig. 1c). Periods of IBE between DT bouts lasted 15.3 ± 2.8 h (n = 2471 IBE periods, Fig. 1e). Moreover, consistent with prior reports^62^, torpor entry occurred preferentially during subjective night (∼ZT17-18) (n = 3,595 torpor bouts, Fig. 1g) and was characterized by a stereotyped nesting behavior with curled posture and absence of gross motor movement, food, or water intake^2,61^. By contrast, arousal from DT exhibited no clear circadian pattern (Fig. 1h), consonant with disrupted circadian clock oscillations at temperatures below _15°C62,63._

### POA activity regulates hamster entry into deep torpor

Having established laboratory conditions for efficient hibernation induction, we employed this system to investigate neural activity changes associated with DT onset. To this end, we collected brain samples during DT and IBE, as well as from non-hibernating control animals, and performed immunostaining for FOS, a marker of neuronal activity (Fig. 2a). Given prior reports of neuronal activity marker induction in hypothalamic regions during DT bouts^39,62,64^, and classic studies showing that hypothalamic lesions disrupt T_b_ regulation during hibernation^65^, we focused our analysis primarily on the hypothalamus. Among the 32 distinct regions analyzed, we observed DT-active regions associated with circadian rhythm modulation (Suprachiasmatic nucleus)^66–68^, energy homeostasis (Arcuate nucleus)^69–71^, and arousal (Paraventricular nucleus of the thalamus)^72,73^. Notably, the anteroventral periventricular nucleus (AVPe) within the anterior POA exhibited the highest degree of FOS induction in DT relative to IBE or non-hibernating control samples (14.1% vs. 6.9% and 0.8%, P = 0.0685 and P < 0.001, respectively) (Fig. 2b-d, Supplementary Tables 1, 2, Extended Data Fig. 2a, b).

**Fig. 2:**
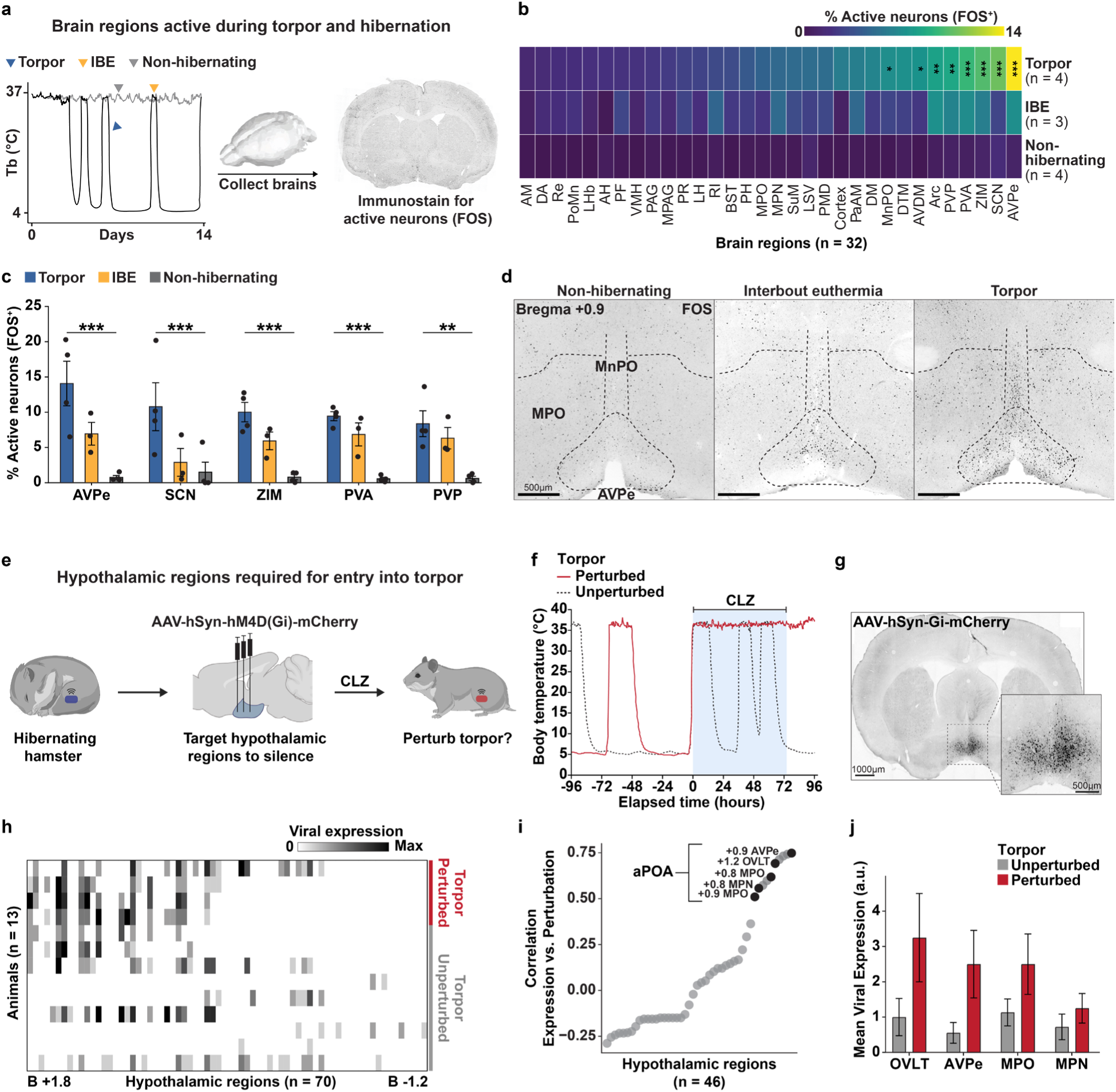
Identification of anterior preoptic area activity as necessary for deep torpor entry. **a**, Strategy for identifying brain regions with hibernation state-dependent activity. Arrows indicate relative sample collection points for torpor, IBE, and non-hibernating animals. **b,c**, Hypothalamic and thalamic regions demonstrating an increase in FOS-positive cells in torpor as compared to both IBE and non-hibernating controls (*P-adjusted < 0.05, **P-adjusted < 0.01, ***P-adjusted < 0.001). **d**, FOS immunostaining in the hamster POA across hibernation states. AVPe, MPO, and MnPO denoted by dashed lines. **e**, Strategy for identifying hypothalamic subregions regulating deep torpor entry. **f**, T_b_ traces showing CLZ-mediated disruption of DT in one Gi-DREADD-mCherry-transduced hamster (solid red) but not the other (dashed black). **g**, Coronal section and inset showing Gi-mCherry expression in the hypothalamus. **h**, Quantification of regional viral Gi-mCherry expression across injected animals. Hypothalamic regions (n = 70) plotted relative to bregma anterior-posterior coordinates. **i**, Correlation of regional viral expression across animals identifies the anterior POA (aPOA) as the region whose inhibition most strongly disrupts DT re-entry. **j**, Higher viral Gi-mCherry expression in aPOA regions from animals exhibiting CLZ-dependent torpor disruption (n = 4) compared to unaffected animals (n = 13). Mean ± s.e.m.

To test directly whether hypothalamic neuronal activity is required for the initiation and maintenance of hibernation, we stereotaxically injected adeno-associated virus (AAV) expressing an inhibitory Gi-coupled DREADD (Designer Receptor Exclusively Activated by Designer Drug; AAV-hSyn-hM4D(Gi)-mCherry)^74^ into hibernating hamsters (Fig. 2e), systematically targeting different areas of the anterior hypothalamus that exhibited substantial DT-associated activity. After allowing animals to recover and resume DT bouts, we administered the DREADD agonist clozapine (CLZ) via drinking water. As animals consumed CLZ during their IBE periods, we observed a cessation of torpor bouts in a subset (4/13) of the tested animals, while all animals that underwent sham surgery and received CLZ (5/5) readily re-entered torpor (Fig. 2f-h, Extended Data Fig. 2c). Correlation of the pattern of Gi-DREADD expression across animals with the degree of DT perturbation identified subregions of the anterior POA (aPOA), including the medial preoptic area (MPO), medial preoptic nucleus (MPN), and AVPe, as regions whose silencing was associated with perturbed hibernation (Fig. 2h-j, Supplementary Table 3). Together, these findings implicate the anterior preoptic area as a key region whose activity is necessary for the initiation of DT bouts during hibernation.

### Cross-species transcriptomic profiling identifies DT-active neuronal populations

The aPOA has been implicated in the regulation of fasting-induced daily torpor (FIT) in mice^39–44^. Our chemogenetic silencing studies in hamster therefore raised the possibility that this region plays a conserved role in the control of hypometabolic states across mammals. To gain further insight into the diversity of neural cell types in the hamster POA and their relation to the cellular composition of the murine POA, we dissected and profiled the POA of 7 adult hamsters (4 females and 3 males) using single-nucleus paired RNA- and ATAC-sequencing (multiome) (Fig. 3a). We annotated the hamster POA cells by aligning and integrating the resulting data with recently published mouse POA multiome and spatial transcriptomic datasets^75,76^, generating the first multiomic cross-species POA cellular taxonomy (Extended Data Fig. 3-5).

**Fig. 3:**
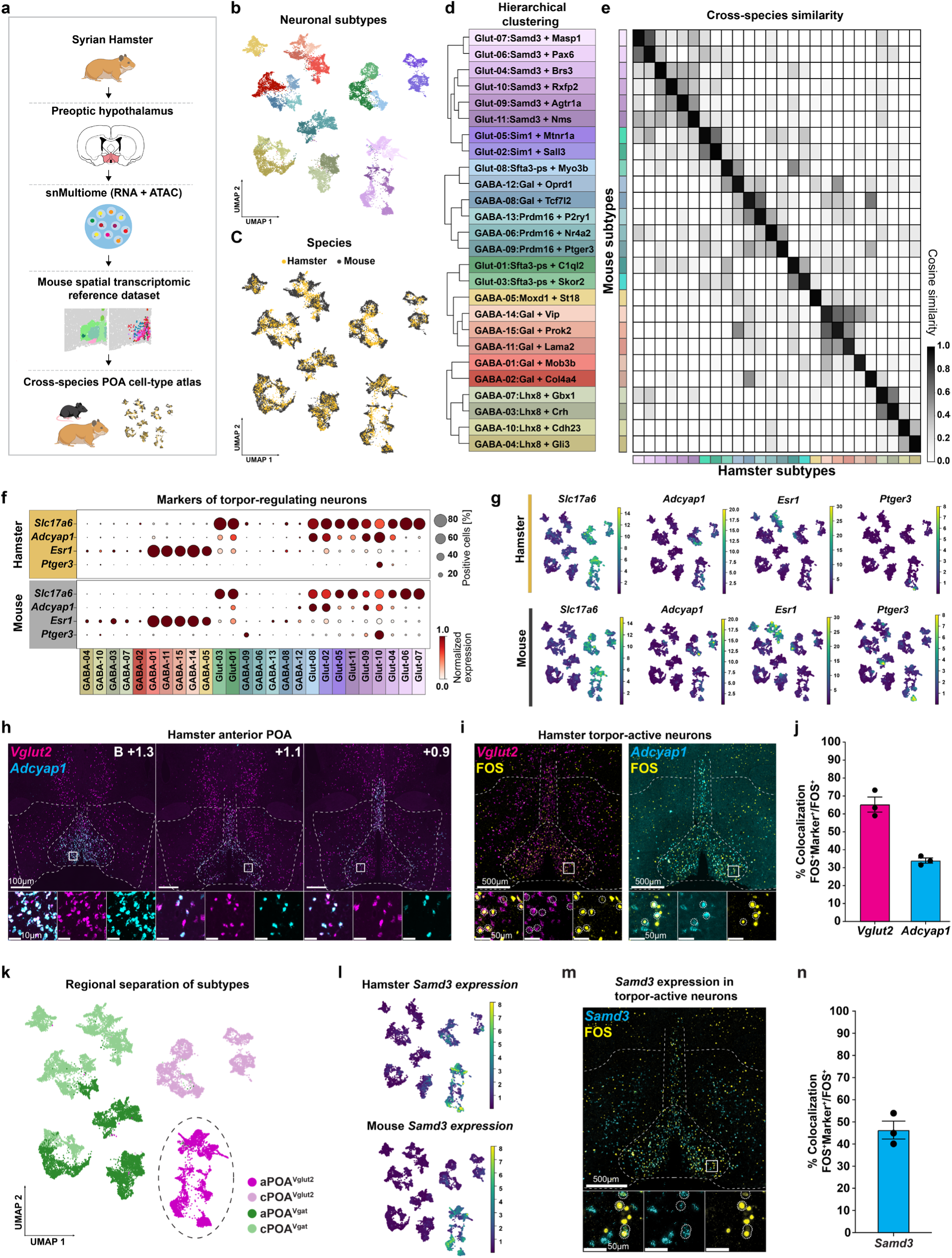
Cross-species multi-omic mapping identifies a glutamatergic aPOA neuronal population active during deep torpor. **a**, Workflow to generate a cross-species POA cellular taxonomy. **b,c**, Uniform manifold approximation and projection (UMAP) plot showing 18,406 nuclei from mice (n = 11) and 8,355 nuclei from Syrian Hamsters (n = 7), indicating 15 GABAergic and 11 glutamatergic neuronal subtypes (**b**) or species of origin (**c**). **d**, Hierarchical clustering of the neuronal subtypes. **e**, Cosine similarity analysis showing pairwise transcriptional relatedness between hamster and mouse neuronal subtypes based on mean gene expression. **f**, Dot plot showing FIT-associated marker gene expression across mouse and hamster neuronal subtypes. **g**, UMAP feature plots depicting marker gene expression in the indicated species. **h**, ISH for select FIT-associated marker genes in hamster aPOA. **i,j**, Combined FOS immunostaining and ISH analysis for *Slc17a6* (*Vglut2*) and *Adcyap1* from aPOA of hamsters (n = 3) in DT, with accompanying quantification. Mean ± s.e.m. **k**, UMAP plot annotating hamster and mouse POA neurons by regional location based on mouse MERFISH data. **l**, UMAP feature plots depicting *Samd3* expression in the indicated species. **m,n**, Combined FOS immunostaining and *Samd3* ISH analysis of aPOA from hamsters (n = 3) in DT, with accompanying quantification. Mean ± s.e.m.

Since activity of brain regions within the median and medial POA was essential for DT entry (Fig. 2i), we focused our analysis on these regions. Graph-based clustering identified major neuronal and non-neuronal cell classes, with further subclustering of neuronal cells (*n* = 26,761 cells) identifying 15 distinct GABAergic and 11 glutamatergic neuronal cell types shared between hamster and mouse (Fig. 3b-d, Extended Data Fig. 4). Importantly, every neuronal cluster contained both mouse and hamster cells (Fig. 3c), and pair-wise transcriptional similarity analysis (cosine similarity score) between all hamster and murine neuronal cell clusters revealed consistent 1:1 cross-species relationships, further supporting the validity of our cross-species atlas alignment (Fig. 3e). Moreover, with the exception of a few cell types, the relative cell type abundance across POA neuronal clusters was largely preserved between hamster and mouse (Extended Data Fig. 5e). Thus, our analyses reveal broad cellular homology between hamster and mouse POA.

Focusing more closely on POA neuronal populations implicated in FIT in the mouse, we found that the expression patterns of molecular markers of mouse FIT-associated neurons (*Slc17a6/Vglut2, Adcyap1, Esr1,* and *Ptger3*) were broadly conserved across hamster and mouse POA cell types (Fig. 3f, g). Moreover, *in situ* hybridization (ISH) analysis indicated that *Vglut2* and *Adcyap1*, two broad markers of FIT-regulating cells, labeled neurons in similar anatomical locations in the mouse and hamster aPOA, including the AVPe, MPO, and MnPO (animals = 4, Fig. 3h, Extended Data Fig. 6). Leveraging these findings, we carried out combined immunostaining and ISH on aPOA samples from hamsters (n = 3) in deep torpor and found that the majority of DT-active (FOS-positive) neurons express *Slc17a6/Vglut2* (65.2 ± 4.2%, animals = 3), with a considerable portion expressing *Adcyap1* (34.0 ± 1.4%, animals = 3), and relatively few expressing *Slc32a1/Vgat* (12.8 ± 1.5%, animals = 3, Fig. 3i, j, Extended Data Fig. 7a, b). Together, these findings suggest that homologous aPOA glutamatergic neuronal populations may be active during mouse FIT and hamster DT.

To further test this idea, we leveraged anatomical information from our transcriptomic and anatomical atlas to identify Glut-04, -06, -07, -09, -10, and -11 clusters as corresponding to FIT-regulating neurons (hereafter collectively termed aPOA^Vglut2^ neurons; Fig. 3k, Extended Data Fig. 7c), and expression of the *Samd3* gene was found to exclusively mark this population (Fig. 3l). Indeed, a substantial fraction of hamster DT-active (FOS+) neurons also expressed *Samd3* (46.4 ± 4.1% of FOS+ AVPe neurons are positive for *Samd3* mRNA, Fig. 3m, n), further supporting the idea that aPOA^Vglut2^ neurons are active during DT entry and might regulate entry into this state. Together, our findings establish the first cross-species atlas of the aPOA, a key region whose activity is necessary for the initiation of DT bouts during hibernation, uncovering broad homology between mouse and hamster cell types and implicating aPOA^Vglut2^ neurons as promising candidate drivers of DT entry in the hamster.

### Enhancer-driven viral tools to access aPOA^Vglut2^ neurons

To assess the functional role of aPOA^Vglut2^ neurons in hamster DT entry, we required the ability to selectively target and manipulate these cell populations *in vivo*. Transgenic options in the hamster remain limited, and a *CamkIIa* promoter-driven AAV exhibits poor specificity for glutamatergic neurons in the hamster POA (42.6 ± 2.0%, animals = 4, Extended Data Fig. 8). Recently, AAV constructs incorporating compact cell-type-specific enhancer gene regulatory elements (GREs) have offered a promising alternative to traditional transgenic technologies in nonstandard species^77–83^. We therefore sought to generate new enhancer-AAV tools driving restricted payload expression in the target aPOA^Vglut2^ cell population.

To develop a viral approach to selectively target aPOA^Vglut2^ neurons, we leveraged the extensive murine single-nucleus chromatin accessibility (snATAC-seq) data included in our multiomic cross-species atlas to identify conserved genomic elements exhibiting selectively enhanced accessibility in aPOA^Vglut2^ neuron cell populations (Fig. 4a). First, we employed a modeling-based approach to identify cell-type-specific nucleosome-depleted gene regulatory regions. Subsequently, we performed differential accessibility analysis to identify putative GREs with enriched accessibility in aPOA^Vglut2^ neuronal populations relative to other neuronal subtypes (Fig. 4b, Extended Data Fig. 9), and filtered GREs based on proximity to aPOA^Vglut2^ neuron-restricted marker genes (Fig. 4c).

**Fig. 4:**
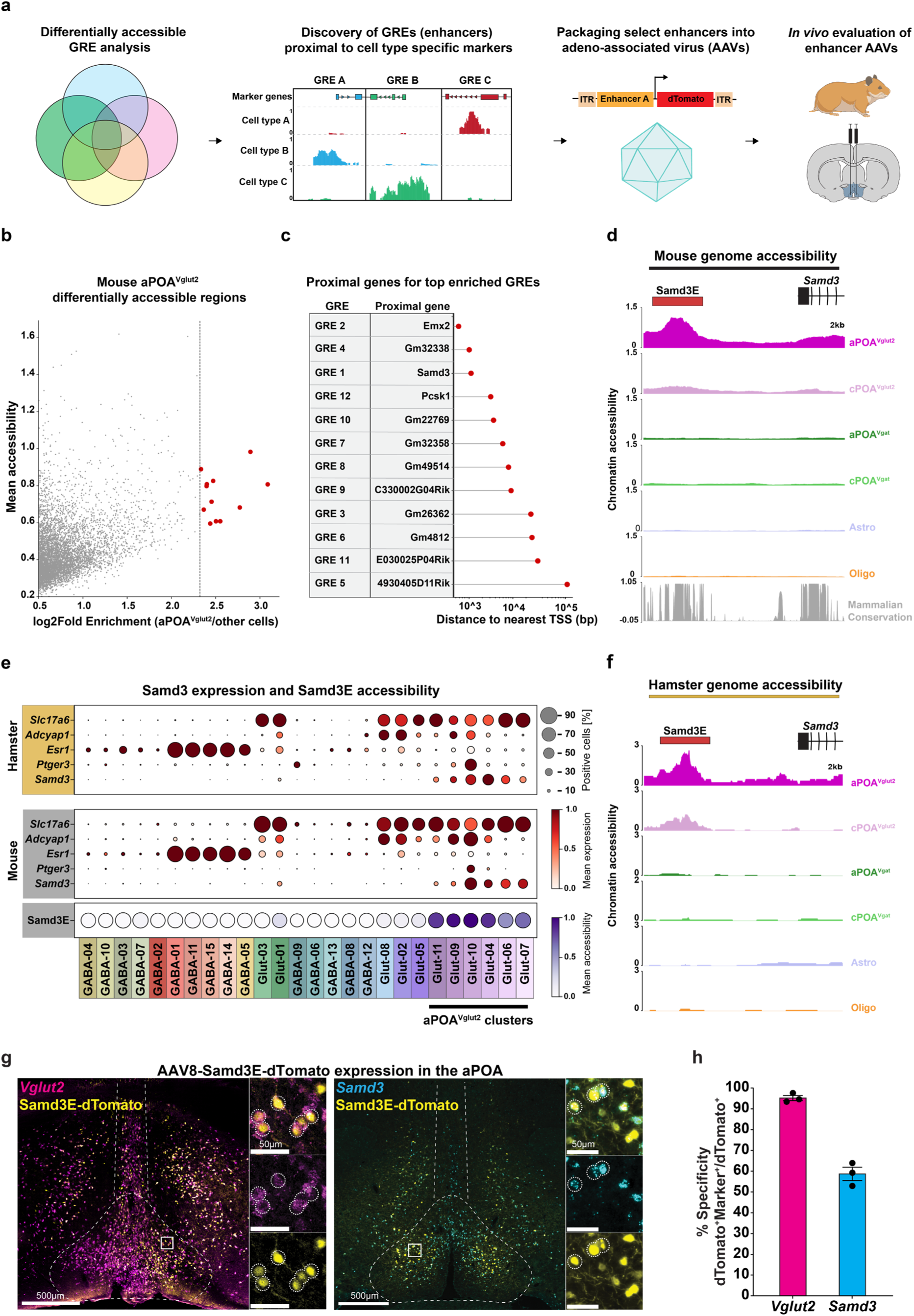
An enhancer AAV enabling selective targeting of aPOA^Vglut2^ neurons in hamsters. **a**, Workflow for generation of aPOA^Vglut2^ neuron-selective enhancer-AAVs. **b**, Scatter plot of genomic elements (n = 15,208) based on differential accessibility in aPOA^Vglut2^ neurons and mean aPOA^Vglut2^ accessibility signal. Top 10 most differentially accessible elements (FDR-corrected q < 10^-300^) highlighted in red. **c**, Distance to proximal gene transcription start site for the top 10 putative GREs from (**b**). **d**, ATAC-seq genome browser tracks (normalized counts per location) for the mouse *Samd3*-proximal putative GRE (*Samd3E,* red box) in the indicated cell populations in mouse, with accompanying sequence conservation below. **e**, Dotplot depicting *Samd3* gene expression across mouse and hamster POA neuronal subtypes, along with *Samd3E* accessibility in the mouse. **f**, ATAC-seq genome browser tracks (normalized counts per location) for the hamster *Samd3*-proximal putative GRE (*Samd3E*, red box) in the indicated cell populations in hamster. **g,h**, Combined immunostaining for viral dTomato and ISH analysis for *Slc17a6* (*Vglut2*) and *Samd3* from aPOA of hamsters injected with AAV8-Samd3E-dTomato, with accompanying quantification. Mean ± s.e.m. (n = 3 animals).

Notably, using this approach, we identified a conserved 500 bp element proximal to the mouse *Samd3* gene, hereafter termed *Samd3E*, that exhibited the highest accessibility enrichment in aPOA^Vglut2^ neurons relative to caudal POA^Vglut2^ neurons and POA^Vgat^ neurons (FDR < 10^-300^, log_2_Fold Enrichment = 3.1, Fig. 4b-e), with similar aPOA^Vglut2^ neuron-enriched accessibility also observed in the hamster (P = 10^-173^, log_2_Fold Enrichment versus other populations = 3.2, Fig. 4f, Extended Data Fig. 9).

The resulting candidate mouse GRE sequences were synthesized and cloned into an AAV backbone upstream of a minimal beta-globin promoter sequence and a red fluorescent protein (RFP) reporter transgene prior to packaging into functional AAV8 particles for *in vivo* evaluation. Notably, the *Samd3E* element conferred remarkable specificity, restricting reporter expression almost exclusively to *Vglut2*+ neurons in the hamster aPOA (95.2 ± 1.2%; animals = 3), with substantial co-labeling of *Samd3+* cells (58.7 ± 3.2%, animals = 3, Fig. 4g, h, Extended Data Fig. 10). Taken together, these findings highlight the power of this approach to generate viral tools to study brain circuits across species, including in hibernating animals, and establish the *Samd3E*-driven AAV (Samd3E-AAV) as an effective tool for achieving targeted genetic access to glutamatergic neurons of the anterior POA.

### aPOA^Vglut2^ neuronal activity is necessary and sufficient for deep torpor entry

Employing our newly developed Samd3E-AAV, we set out to address the role of aPOA^Vglut2^ neuronal activity in hibernation and DT by expressing an inhibitory Gi-DREADD (AAV-Samd3E-hM4D(Gi)-EGFP) in hamster aPOA^Vglut2^ neurons (Fig. 5a). Neuronal inhibition via CLZ administration had no significant effect on basal homeostatic T_b_ or circadian T_b_ rhythmicity in control non-hibernating Gi-DREADD-expressing hamsters (Extended Data Fig. 11a, b). However, actively hibernating Gi-DREADD-expressing (n = 12) animals showed a significant delay in the resumption of DT bouts following CLZ treatment compared to when these same animals were administered a water-only vehicle control (114.9 ± 45.4 hours vs. 64.1 ± 24.5 hours, P = 0.0003). Indeed, a majority (7/12) of CLZ-treated Gi-DREADD-expressing animals failed to re-enter DT during the full 6-day course of CLZ treatment, whereas none (0/12) of the animals failed to resume DT during treatment with the vehicle control. Moreover, animals (n = 5) expressing a control reporter payload (AAV-Samd3E-EGFP-Synaptophysin-mRuby) exhibited no significant difference in the time to DT resumption following CLZ or vehicle treatment (61.6 ± 26.5 hours vs. 47.4 ± 10.2 hours, respectively. P = 0.6167), with all animals resuming torpor bouts within the CLZ treatment window (Fig. 5b, c). Together, these results strongly suggest that aPOA^Vglut2^ neuronal activity is required for entry into hibernation-associated DT.

**Fig. 5:**
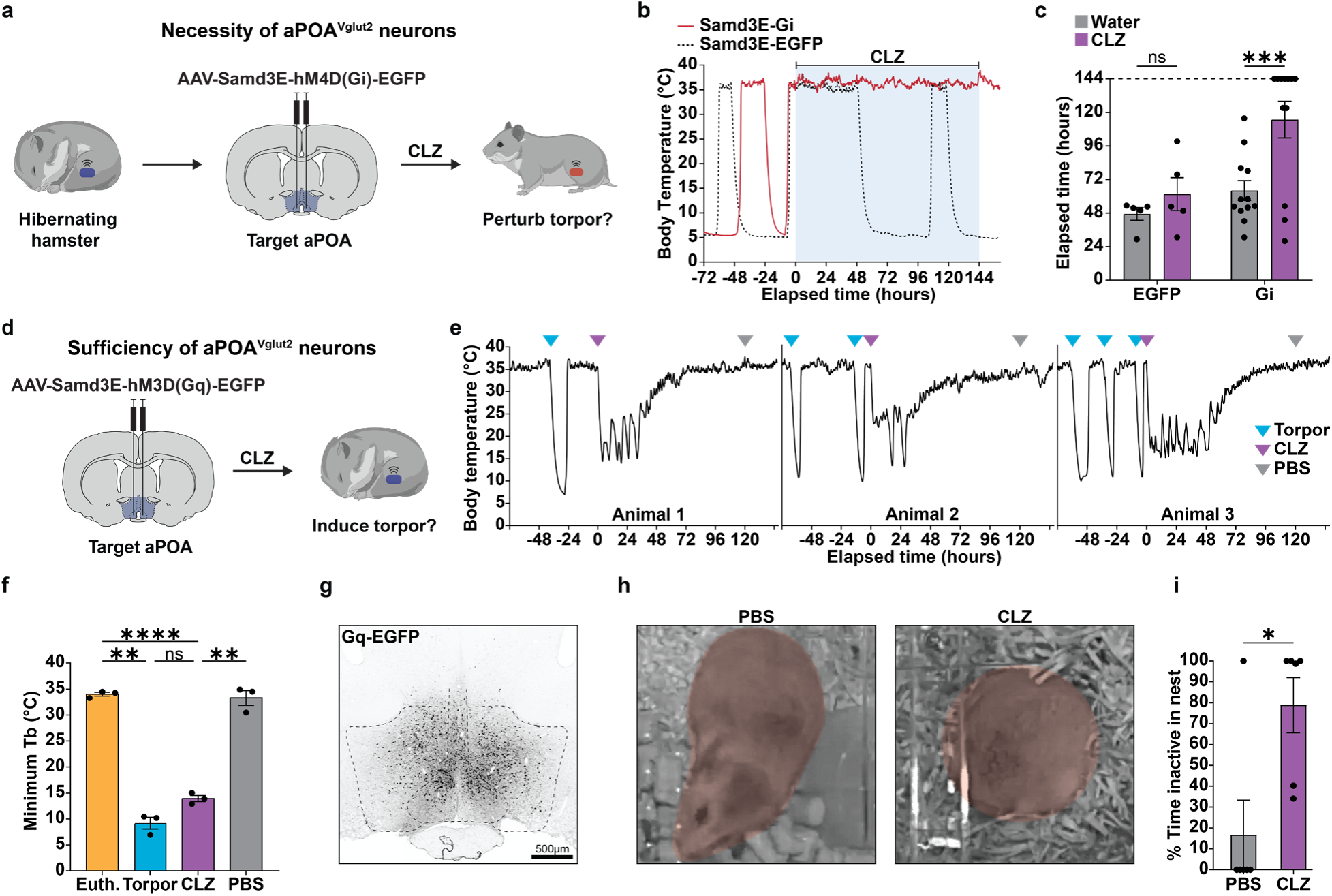
aPOA glutamatergic neuron activity is necessary and sufficient to induce deep torpor-like entry in hamsters. **a**, Strategy for testing the necessity of aPOA^Vglut2^ neuronal activity for DT reentry. **b**, T_b_ traces demonstrating perturbation of torpor resumption upon CLZ administration in Samd3E-Gi-DREADD-but not control Samd3E-EGFP-transduced hamsters. **c**, Samd3E-Gi-DREADD-transduced hamsters (n = 12), but not control Samd3E-EGFP-transduced animals (n = 5), show significant perturbations in torpor initiation when administered with CLZ but not water (Šídák’s multiple comparisons test). **d**, Strategy for testing the sufficiency of aPOA^Vglut2^ neuronal activity for DT entry. **e**, T_b_ traces from Samd3E-Gq-DREADD-injected animals illustrating natural torpor bouts as well as artificial bouts elicited by CLZ administration. **f**, Quantification of minimum T_b_ reached in Samd3E-Gq-DREADD-transduced animals during euthermic periods, natural deep torpor, and following administration of CLZ or control PBS (n = 3, Tukey’s multiple comparisons test). **g**, Coronal section showing Samd3E-Gq-DREADD-EGFP expression in the hamster aPOA. **h**, Images of a Samd3E-Gq-DREADD-transduced hamster following PBS or CLZ treatment. **i**, Percentage of time Samd3E-Gq-DREADD-transduced animals (n = 6) spent inactive in the nest over a 10 min observation window CLZ or control PBS administration (paired t-test). ****P-adjusted < 0.0001, ***P-adjusted < 0.001, **P-adjusted < 0.01, *P-adjusted < 0.05, ns = not significant).

To determine the extent to which aPOA^Vglut2^ neuron activity is sufficient to drive DT entry, we selectively introduced an excitatory Gq-DREADD into hamster aPOA^Vglut2^ neurons via our Samd3E-AAV (AAV-Samd3E-hM3D(Gq)-EGFP, Fig. 5d). CLZ administration led to a rapid, significant drop in T_b_ in Gq-DREADD-expressing hamsters, with the animals reaching a minimum body temperature of 13.9 ± 1.0°C (P < 0.0001 relative to normothermia) and maintaining a reduced body temperature for 56.9 ± 13.1 hours, consistent with the induction of a prolonged torpor-like state (Fig. 5e-g). By contrast, these same animals exhibited no discernable T_b_ change upon control PBS administration (minimal T_b_ = 33.3 ± 2.4°C, P = 0.8959 relative to normothermia). In addition, CLZ treatment of Gq-DREADD-expressing animals (n = 6), but not PBS treatment of the same animals (n = 6) or PBS/CLZ treatment of hamsters injected with a control viral construct (n = 3), induced behaviors consonant with torpor entry, including a curled posture and prolonged inactivity in the nest (P = 0.0168, Fig. 5h, i, Extended Data Fig. 12, Supplementary Video 1). Moreover, follow-up analyses revealed that the depth of CLZ-induced torpor across animals correlated with levels of aPOA Gq-DREADD expression, suggesting that more extensive aPOA^Vglut2^ neuron activation drives deeper torpor-like bouts (r = -0.9577, P = 0.0104, Extended Data Fig. 11c, d). Together, these findings show that aPOA^Vglut2^ neuronal activity is both necessary and sufficient to initiate a profound T_b_ decrease and DT-associated nesting behaviors in the hamster.

Finally, given that aPOA^Vglut2^ neurons are present in both the Syrian hamster (a hibernator) and mouse (non-hibernator), we investigated whether virus-mediated chemogenetic activation of this neuronal population in mice housed at low ambient temperatures would drive a DT-like decrease in T_b_. Indeed, CLZ treatment of Samd3E-Gq-DREADD-transduced mice yielded a rapid and sustained drop in body temperature, although markedly less than that observed in hamsters (Extended Data Fig. 13). Thus, aPOA^Vglut2^ neuronal activation is sufficient to elicit acute, pronounced reductions in core temperature in both hibernating and non-hibernating species, with the difference in magnitude suggestive of potential hibernator-specific evolutionary modifications in downstream effector circuitry.

## Discussion

Hibernation is a recurrently evolved survival strategy across the mammalian lineage defined by a regulated multi-day decrease in metabolic rate and body temperature, known as torpor^2,3^. Entry into this state is orchestrated by a complex interplay of environmental cues and internal signals^84^. However, despite decades of work, including reports of hibernation-associated neural activity changes and classic lesion studies implicating the hypothalamus^39,62,64,65^, the specific neuronal populations that initiate entry into hibernation-associated torpor have remained undefined. Here, using a Syrian hamster model, we combine neural activity mapping with chemogenetic manipulation to demonstrate that activity within the aPOA is required for entry into deep torpor entry.

Our single-nucleus transcriptional and epigenomic profiling of the hamster POA revealed strong cellular homology with the mouse, and surprisingly no evidence of hibernator-specific neuronal cell types. Instead, we observed that *Samd3-*expressing aPOA glutamatergic neurons exhibit elevated activity upon hibernation-associated DT in the hamster. Leveraging our multiomic POA atlas, we developed an enhancer-driven AAV that enables selective access to this population. Using this tool, we show that silencing aPOA^Vglut2^ neurons blocks deep torpor entry, whereas chemogenetic activation of the same population is sufficient to drive a rapid reduction in body temperature accompanied by torpor-associated nesting behavior. Together, these studies establish Samd3E-AAV-targeted aPOA^Vglut2^ neurons as key regulators of hibernation entry and provide the first genetic access to neural circuits controlling this complex physiological state. Moreover, the Samd3E-AAV and the underlying enhancer-driven viral targeting strategy will be broadly applicable for dissecting hibernation circuit elements in other species and examining whether hibernation circuitry is conserved across hibernators, including primates.

Although fasting-induced daily torpor and seasonal hibernation both involve reductions in body temperature, metabolic rate, and heart rate, these states differ markedly in their depth, duration, and ethological context. FIT in mice represents a short-term response to food scarcity, lasting only hours^2,3,85–87^, whereas hibernation is a seasonal behavior involving deeper, multi-day torpor bouts punctuated by brief periods of interbout euthermia. While the evolutionary origin of these adaptations remains a subject of debate, our findings raise the possibility that differences between daily torpor and hibernation arise from species-specific connectivity or activity patterns within a broadly conserved aPOA^Vglut2^ neuronal population. These differences are potentially shaped by cross-species gene expression differences we observe in these cells (Extended Data Fig. 7d). Consistent with this idea, chemogenetic activation of aPOA^Vglut2^ neurons in hamsters elicits substantially deeper torpor-like bouts than in mice. It is notable that chemogenetically-induced torpor approached but did not fully recapitulate the body temperature observed in natural deep torpor. This difference may reflect lower viral transduction/targeting efficiency (Extended Data Fig. 10) or may indicate additional circuit components required for full expression of deep torpor. Future studies will be needed to determine whether this aPOA^Vglut2^ population or the broader hibernation circuitry undergoes active remodeling during the prolonged pre-hibernation conditioning period.

More broadly, our findings of a shared circuit node regulating daily torpor and seasonal hibernation provide further support for the idea that regulated heterothermy is an ancestral mammalian trait^88,89^. This would further suggest that core elements of torpor-inducing circuitry are conserved across all mammals, including in species that do not naturally hibernate. Consistent with this notion, *in situ* hybridization data from the marmoset (*Callithrix jacchus*), a non-hibernating primate, indicate that neurons expressing torpor-associated markers are present in homologous regions of the aPOA^90–92^ (Extended Data Fig. 14). The Samd3E-AAV that we developed is an ideally suited tool to investigate the degree to which activating these torpor-associated neuronal population in non-hibernators, including non-human primates, could induce a synthetic torpor-like state.

Together, we identify a neuronal population that controls hibernation, enabling for the first time the ability to initiate, manipulate and study this fascinating adaptation. Our work establishes a foundation for future efforts to define the broader neural architecture of hibernation, including upstream pathways conveying seasonal and metabolic cues, downstream effectors mediating systemic adaptation, and the cellular mechanisms that permit neurons and other organs to function at near-freezing temperatures. Beyond advancing our understanding of mammalian physiology, these insights may ultimately inform approaches to harness hibernation-related mechanisms for therapeutic hypothermia, organ preservation, metabolic control, protection from muscle atrophy, aging-related decline, and perhaps even space travel.

## Methods

### Animals

Animal experiments were approved by the Massachusetts Institute of Technology Committee of Animal Care (Protocol 2503000788), following ethical guidelines described in the US National Institutes of Health Guide for the Care and Use of Laboratory Animals and the Animal Welfare Act.

#### Hamster

For all experiments, unless otherwise noted, adult female (5-6-week-old) Syrian hamsters (Inotiv, Strain HsdHan:AURA, Stock 089) were used. Hamsters were singly caged and initially housed at 22°C under a standard 12-hour light/dark cycle (12L:12D) cycle in temperature- and humidity-controlled rooms and had *ad libitum* access to food and water.

#### Mouse

Adult female (8-week-old) C57BL/6J (JAX #000664) mice were used. All mice were group housed initially at 22°C under a standard 12L:12D cycle in temperature-and humidity-controlled rooms and had *ad libitum* access to food and water.

No statistical methods were used to predetermine the sample size. Hamsters were randomly assigned to experimental groups before surgery. Where possible, investigators were blinded during analysis.

### Hibernation Induction

Utilizing a modified hibernation paradigm^61,93^ female (5-6-week-old) hamsters were singly caged and housed at 22°C under a standard 12L:12D cycle for a short period to acclimate to the animal facility. Upon completion of acclimation, hamsters were transferred to a reduced 8L:16D cycle with an ambient temperature of 22°C. After 7-8 weeks, hamsters were placed in temperature-controlled chambers with 8L:16D cycle and ambient temperatures of 4-5°C. Initiation of hibernation was confirmed by changes in T_b_ measured by telemetric temperature probes. In all cases, hamsters had *ad libitum* access to food and water.

### Telemetric monitoring of core body temperature

Within the 7-8-week period of housing at 22°C in 8L:16D cycle, hamsters were anesthetized with 3% isoflurane and implanted with telemetric/logger temperature probes (Star-Oddi, DST microRF-T) in the abdominal cavity. Hamsters were allowed to recover for a minimum of three days before additional procedures or experiments were performed. After recovery and at the end of the 7-8-week period, hamsters were moved to temperature-controlled chambers, and core body temperature was monitored in home cages placed onto a radio frequency receiver antenna (Star-Oddi, Antenna & RF-box). Upon euthanasia, probes were removed and complete logged data downloaded. Core body temperature was logged every 20-30 min.

### Torpor bout analysis

#### Classification

To classify torpor bouts, we employed a Gaussian Mixture Model in R. The classification was based on minimum body temperature and bout duration. Each bout was assigned to the cluster in which it had the highest posterior probability of membership. A cluster characterized by lower T_b_ and longer duration was designated as “Deep”, whereas the other cluster was classified as “Shallow”. Furthermore, periods of between bouts above the set T_b_ threshold were classified as IBE.

#### Circadian preference

In R, circadian preference for torpor and IBE initiation was quantified using the Raleigh test of circular uniformity.

#### Exponential decay

In R, elapsed time in torpor for each body temperature reading from torpor bouts classified as deep were determined for the first 30 hours. Next, using the mean of temperatures at each time across animals, a nonlinear model was fit to the data using the minimum temperature within the 30-hour period as a fixed asymptotic value.

### Viral constructs

AAV8-hSyn-hM4D(Gi)-mCherry (Addgene, 50475-AAV8) was obtained from Addgene. AAV8-Samd3E-dTomato-nlsdTomato, AAV8-Samd3E-hM4D(Gi)-P2A-nlsGFP, AAV8-Samd3E-EGFP-P2A-Synaptophysin-mRuby, and AAV8-Samd3E-hM3D(Gq)-P2A-nlsGFP were produced in-house using the method previously described^94^ and titered via digital PCR (QIAGEN, QIAcuity). All viruses were diluted with PBS to a final titer of 1E13 vg/mL before stereotaxic delivery into the hamster brain. For mice, AAV8-Samd3E-hM3D(Gq)-P2A-nlsGFP with a final titer of 5E13 vg/mL was used.

### Stereotaxic viral injection

#### Hamster

Hamsters were anaesthetized with 3% isoflurane and placed in a stereotaxic head frame (David Kopf Instruments, model 1900). If hamsters were actively hibernating, they were aroused from hibernation and placed at 22°C the day prior to surgery. An air-based injection system connected to a pulled glass needle was used to infuse ∼100 nL of virus bilaterally at regions of interest. Hypothalamic regions were targeted by referencing the hamster stereotaxic atlas^95^. When targeting the aPOA, coordinates +1.2 mm AP, ±0.4 mm ML, -7.5mm DV were used. Hamsters were allowed to recover at 22°C for a minimum of 3 days before a return to 4°C chambers. Animals were afforded a recovery period of at least two weeks post-surgery prior to further experimental manipulation to allow for viral expression and, if appropriate, resumption of hibernation.

#### Mouse

Mice were anesthetized with 3% isoflurane and placed in a stereotaxic head frame (RWD, Guangdong, China). A Nanoject III (Drummond Scientific, Broomall, PA) system connected to a pulled glass needle was used to infuse ∼100 nL of virus bilaterally into the aPOA (distance from Bregma, AP: +0.6 mm, ML: ±0.4 mm, DV: -5.2 mm). Mice were monitored for three days post-surgery to ensure full recovery, with a waiting period of at least two weeks to allow sufficient viral expression.

### Chemogenetic manipulations

#### Hamster

Clozapine dihydrochloride (Hello Bio, HB6129) was either dissolved in sterile water or PBS at a concentration of 10 mg/mL and then stored at -20°C until use. For administration via drinking water, CLZ dissolved in sterile water was diluted in drinking water at a dose of 0.1 mL/kg (1 mg/kg) in a final volume of 250 mL. Animals were maintained on CLZ water for 3-7 days. For intraperitoneal (I.P.) administration, CLZ dissolved in PBS was administered at a dose of 0.5 mL/kg (5 mg/kg). In each case when treating actively hibernating hamsters, they were aroused from hibernation the morning of each experiment and treatment was administered in the evening.

#### Mouse

Mice injected with AAV8-Samd3E-hM3D(Gq)-P2A-nlsGFP were single housed and placed in a temperature-controlled chamber at 8°C for three days. After three days, mice were I.P. injected with CLZ at a dose of 5 mg/kg and surface body temperature was measured using a thermal camera (Teledyne FLIR C5, Teledyne Flir, Wilsonville, OR). Images were taken before CLZ injection, and 30 min, 60 min, 3 hours, 6 hours and 24 hours after CLZ injection. Analysis of surface body temperature and representative images were generated using the Flir Ignite software (https://ignite.flir.com/).

### Immunofluorescence

Hamsters were deeply anesthetized with isoflurane and then euthanized by transcardial perfusion of 50 mL ice-cold 1X PBS followed by 50 mL of 4% paraformaldehyde (PFA). Brains were extracted, post-fixed overnight with 4% PFA at 4°C, and then embedded in 1X PBS with 3% agarose. Brains were sliced on a vibratome into 50 μm coronal sections and stored at -20°C in cryoprotectant buffer (30% sucrose, 1% polyvinyl-pyrrolidone (PVP-40), 30% ethylene glycol, in 0.1 M phosphate buffer). Free-floating coronal sections were washed three times with 1X PBS and then blocked for 1 hour at room temperature with 1X PBS containing 10% donkey serum and 0.3% Triton X-100. Sections were incubated overnight at 4°C with primary antibodies (Rabbit anti-FOS, Synaptic Systems, 226-008; Rabbit anti-RFP, Rockland, 600-401-379; Chicken anti-GFP, Aves Labs, GFP-1020) diluted in 1X PBS containing 5% donkey serum and 0.3% Triton X-100, washed again three times with 1X PBS, and incubated for 2 hours at room temperature with secondary antibodies in 1X PBS containing 5% donkey serum and 0.3% Triton X-100. After washing three times in 1X PBS, samples were counter-stained for 15 minutes at room temperature with 1 μg/mL DAPI, washed again three times in 1X PBS, and then mounted onto SuperFrost Plus glass slides and coverslipped using Fluoromount-G.

### Determining deep torpor-active brain regions

Brains were collected in torpor, IBE, and from non-hibernating control animals and were processed as indicated in the section ‘Immunofluorescence’ using the following reagent for the detection of FOS: Rabbit anti-FOS, Synaptic Systems, 226-008.

#### Sample imaging

Sections were imaged with a TissueFAXS SL whole-slide scanning system equipped with a Zeiss Axio Imager Z2 fluorescence microscope and TissueFAXS image acquisition software (Ragon Institute Microscopy Core).

#### Image analysis

Using TissueQuest analysis software (TissueGnostics GmbH, Vienna, Austria), brain regions were manually annotated and nuclear DAPI signal segmented. Segmented cells containing putative FOS signal were determined to be FOS-positive based on diameter and mean fluorescence intensity. In R, regional cell counts were analyzed and percent active neurons determined by the fraction of FOS-positive and total DAPI-positive cells. Significance was calculated using a mixed-effects ANOVA with post-hoc pairwise comparisons. Pairwise comparison p-values were multiple tests corrected using the Bonferonni method.

### Mapping of virally injected hypothalamic nuclei and correlation with chemogenetically induced torpor perturbation

Hibernating hamsters were bilaterally injected with AAV8-hSyn-hM4D(Gi)-mCherry. Upon resumption of hibernation, hamsters were administered CLZ water for ∼3 days (74 hours) to inhibit virally injected regions, and their core body temperature were recorded.

#### Temperature data analysis

Upon CLZ water treatment, hamsters reinitiating deep torpor within 48 hours were classified as unperturbed, and those failing to reinitiate (remaining euthermic) within this time were classified as perturbed. Sham surgery control animals treated with CLZ water for 7 days informed this threshold (Extended Data Fig. 2c). Absolute elapsed time of euthermia from start of treatment was used for further analysis.

#### Sample imaging

Given the unavoidable variability in hypothalamic region targeting and viral spread, all hamsters were euthanized for post hoc assessment of transduced brain regions. Collected brains were processed as indicated in the section ‘Immunofluorescence’ to detect endogenous mCherry fluorescence. Sections were imaged with a TissueFAXS SL whole-slide scanning system equipped with a Zeiss Axio Imager Z2 fluorescence microscope and TissueFAXS image acquisition software (Ragon Institute Microscopy Core).

#### Image and correlation analysis

Viral mCherry expression was quantified across 70 hypothalamic regions in a semi quantitative manner, where the viral expression was scored 0 (none), 1 (minimal), 2 (partial), or 3 (complete) in each region and hemisphere. The analysis was performed blinded to whether perturbation of torpor was observed in each hamster. To assign a single numeric value for each hypothalamic region in each hamster, we added the semiquantitative expression values from each hemisphere. Next, a Pearson correlation was calculated for each region across all hamsters between the viral expression in that region and the length of time spent euthermic.

### Nuclei dissociation and single-cell analysis

Preoptic area tissue was dissected from adult male and female Syrian hamsters and immediately flash-frozen. The 10X genomics nuclei isolation kit (1000448) was used to obtain dissociated nuclei. Tissue was homogenized on ice in a detergent-based nuclei isolation buffer, passed through a 40-µm filter, and purified by centrifugation. Nuclei were washed, and counted before loading. Single-nucleus multiome libraries were prepared using the 10x Genomics Chromium Single Cell Multiome ATAC + Gene Expression workflow following the manufacturer’s protocol. Constructed libraries were quantified, pooled, and sequenced on an Illumina platform to recommended depth for both modalities. The raw data were processed in parallel with cellranger-arc to generate gene-expression count matrices, ATAC fragment files, and peak-by-cell accessibility matrices. A recently published mouse POA multiome dataset generated with the same 10x system was obtained to build a reference dataset. A combination of scVI and Scanpy^96,97^ pipelines were used to process and integrate the two datasets, including quality control (filtering by gene counts, UMI counts, and mitochondrial RNA content), normalization, log-transformation, identification of highly variable genes, dimensionality reduction, neighborhood graph construction, and Leiden clustering.

To ensure that downstream analyses focused on POA-associated neuronal populations, we used the MERFISH dataset as a spatial reference^76^. To this end, mouse spatial transcriptomic data was subset to only include neurons found in the mPOA and co-clustered with mouse neuronal snRNA-seq. Canonical POA markers and spatial annotations were used to confirm that the identified clusters in both species corresponded to POA-localized neuronal subtypes. Subsequently, the reference mouse dataset was used to identify homologous hamster neurons through integration and label transferring with scANVI^98^. Chromatin accessibility matrices were analyzed with Pycistopic^99^ including psuedobulking, peak calling and filtering, identification of co-accessible regions and differential accessibility analysis.

### Double fluorescent *in situ* hybridization and immunofluorescence

Hamsters were deeply anesthetized with isoflurane and then euthanized by transcardial perfusion of 50 mL ice-cold 1X PBS followed by 50 mL of 4% PFA. Brains were extracted, post-fixed overnight with 4% PFA at 4°C, and then incubated in 1X PBS with 30% sucrose at 4°C until fully infiltrated and sunk to the bottom of the chamber (72-96 hours). Brains were embedded in Optimal Cutting Temperature (O.C.T.) compound, frozen on dry ice, and stored at -80°C until use. Brains were sliced on a cryostat (Leica CM3050 S) into 10-30 μm coronal sections, placed in cryoprotectant buffer, and stored at −20°C until use. Free-floating coronal sections were washed three times with 1X PBS, mounted onto Superfrost Plus slides, and then baked for 45 minutes at 60°C. Subsequent steps followed the standard fluorescent *in situ* hybridization protocol for the RNAscope Fluorescent Multiplex V2 Assay using the following probes: Mau-*Samd3*-C1 (1826261), Mau-*Slc17a6*-C2 (1176381), Mau-*Adcyap1*-O1-C3 (1808861), Mau-*Slc32a1*-C4 (1176381).

Upon completion of *in situ* hybridization, samples immediately were processed according to the ACD RNAscope Fluorescent Multiplex V2 Assay Combined with Immunofluorescence manual with the following modifications: Slide-mounted sections were washed twice in 1X TBS with 0.005% Tween-20 (TBST), blocked for 1 hour at room temperature with 1X TBS containing 10% donkey serum, and incubated overnight at 4°C with primary antibodies (Rabbit anti-FOS, Synaptic Systems, 226-008; Rabbit anti-RFP, Rockland, 600-401-379; Chicken anti-GFP, Aves Labs, GFP-1020) diluted in 1X TBS containing 5% donkey serum. Slides were washed three times with TBST, incubated for 2 hours at room temperature with secondary antibodies in 1X TBS containing 5% donkey serum, washed again three times with TBST, counter-stained for 15 minutes at room temperature with 1 μg/mL DAPI, washed three times with 1X TBS, and finally mounted on coverslips using ProLong Gold.

#### Sample imaging

Sections were either imaged with a TissueFAXS SL whole-slide scanning system equipped with a Zeiss Axio Imager Z2 fluorescence microscope and TissueFAXS image acquisition software (Koch Institute Swanson Biotechnology Center/Microscopy Core) or a Leica Stellaris 8 FALCON confocal microscope (Whitehead Institute W.M. Keck Facility for Biological Imaging).

#### Image analysis

Using QuPath software^100^, colocalization analysis of RNAscope and immunofluorescent signals was performed. After manual annotation of regions of interest, the InstanSeg extension^101^ was utilized for cell segmentation, considering each fluorescent signal. Next, segmented cells containing putative RNAscope and immunofluorescent signals were determined to be positive or negative for associated markers based on diameter and mean fluorescence intensity. Subsequent quantification of counts was performed in R, while plotting and statistical analysis was performed in GraphPad Prism software (Boston, MA).

### Inhibition of aPOA^Vglut2^ neurons

Hibernating hamsters were bilaterally injected with either AAV8-Samd3E-hM4D(Gi)-P2A-nlsGFP or AAV8-Samd3E-EGFP-P2A-Synaptophysin-mRuby. On the day of treatment, hamsters were aroused in the morning, I.P. injected with CLZ and provided with CLZ water in the evening, I.P. injected again 16 hours later, and core body temperature recorded. This regimen accounted for variability in water intake within the first 24 hours of the experiment and maximized inhibition of our targeted population. A CLZ water treatment window of 6 days (144 hours) was used. For PBS control treatment, an identical regimen was followed.

#### Temperature data analysis

In GraphPad Prism (Boston, MA), absolute elapsed time of euthermia from start of treatment was used to determine differences within groups. The treatment window duration (144 hours) was applied as an upper bound for statistical analysis. In R, temperature data from each virus group and treatment condition was mapped to circadian time. In GraphPad Prism (Boston, MA), statistical analysis was performed on temperatures in the light and dark cycles within virus groups.

### Excitation of aPOA^Vglut2^ neurons

Hamsters were bilaterally injected with either AAV8-Samd3E-hM3D(Gq)-P2A-nlsGFP or AAV8-Samd3E-EGFP-P2A-Synaptophysin-mRuby and housed at 4°C to acclimate. On the day of treatment, hamsters were aroused in the morning, I.P. injected with CLZ in the evening, and core body temperature recorded. For PBS control treatment, an identical regimen was followed.

#### Temperature data analysis

In R, temperature data was analyzed using the method described in section ‘Torpor bout analysis’ to determine if the CLZ induced drop in body temperature clustered with either deep or shallow torpor. In GraphPad Prism (Boston, MA), statistical analysis was performed on minimum temperatures across each condition (euthermia, torpor, CLZ, and PBS).

#### Sample imaging

Collected brains were processed as indicated in the section ‘Immunofluorescence’ using the following reagent for the detection of AAV8-Samd3E-hM3D(Gq)-P2A-nlsGFP: Chicken anti-GFP, Aves Labs, GFP-1020. Sections were imaged with a TissueFAXS SL whole-slide scanning system equipped with a Zeiss Axio Imager Z2 fluorescence microscope and TissueFAXS image acquisition software (Koch Institute Swanson Biotechnology Center/Microscopy Core).

#### Image and correlation analysis

aPOA GFP expression was quantified in a semi quantitative manner where expression was scored 0 (none), 1 (minimal), 2 (partial), or 3 (complete) in each hemisphere. The analysis was performed blinded to change in body temperature observed in each hamster. Hemispheric values were summed within animals. In GraphPad Prism (Boston, MA), a Pearson correlation was performed between GFP viral expression and the minimum body temperature.

### Behavioral analysis

Hamsters housed at 4°C and injected with AAV8-Samd3E-hM3D(Gq)-P2A-nlsGFP or AAV8-Samd3E-EGFP-P2A-Synaptophysin-mRuby were I.P. injected with PBS (0.5 mL/kg) after a one-hour baseline video recording of homecage behavior and an additional hour recording post injection. Following at least a week recovery in 4°C chambers, animals were injected with CLZ (5 mg/kg) and videos were recorded for one hour before and post drug administration in 4°C chambers. Homecage behaviors were manually coded using BORIS software^102^. Behavior was coded 30 min after PBS or CLZ injection over a 10 min window.

## Acknowledgments

We thank M. Alkire, M.M. Chen, J.A. Cheng, and N.T. Nigrin for their assistance with mouse experiments. We also thank additional Hrvatin lab members for their thoughtful input on this project and manuscript. The snRNA-seq and snATAC-seq were performed with the Whitehead Institute Genome Technology Core. Imaging was performed either at the Ragon Institute Microscopy Core, Koch Institute Swanson Biotechnology Center/Microscopy Core, or the Whitehead Institute W.M. Keck Facility for Biological Imaging. This work was supported by the Mathers Foundation Grant, Searle Scholars Program, Pew Charitable Trust, Longevity Impetus Grant from Norn Group, Hevolution Foundation and Rosenkranz Foundation, MIT Research Support Committee Grant, McKnight Endowment Fund, and Howard Hughes Medical Institute Hrabowski Scholar Program.

## Author Contributions

A.J.M. and S.H. conceptualized the study and designed experiments. A.J.M, C.M.R, A.J.L.P, W.L., A.S.L. performed and analyzed experiments. E.C.G. and S.H. advised on the study and obtained funding for the research. A.J.M., E.C.G., and S.H. wrote the manuscript.

## Data and materials availability

Data will be made available on GEO. All other data needed to evaluate the conclusions are available in the main text, the extended data, supplementary materials, and available upon request.

## Extended Data

**Extended Data Fig. 1.**
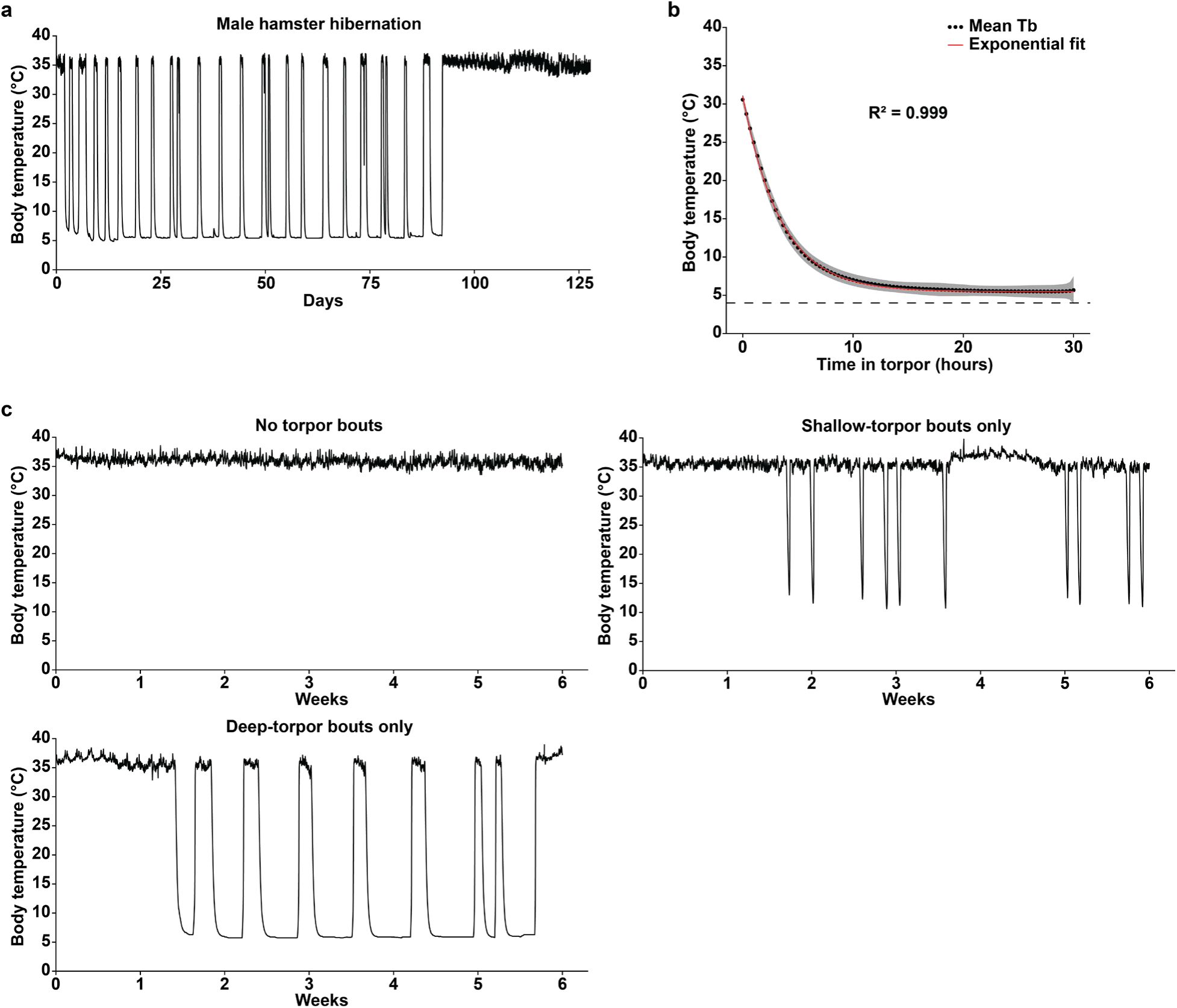
Core body temperature dynamics during hibernation. **a**, Representative core body temperature trace from a male hamster across its hibernation season. **b**, Initial T_b_ decreases during torpor onset follow an exponential decay (animals = 188, torpor bouts = 1262). **c**, Body temperature traces from three animals within a 6-week observation period, demonstrating the range of T_b_ changes.

**Extended Data Fig. 2.**
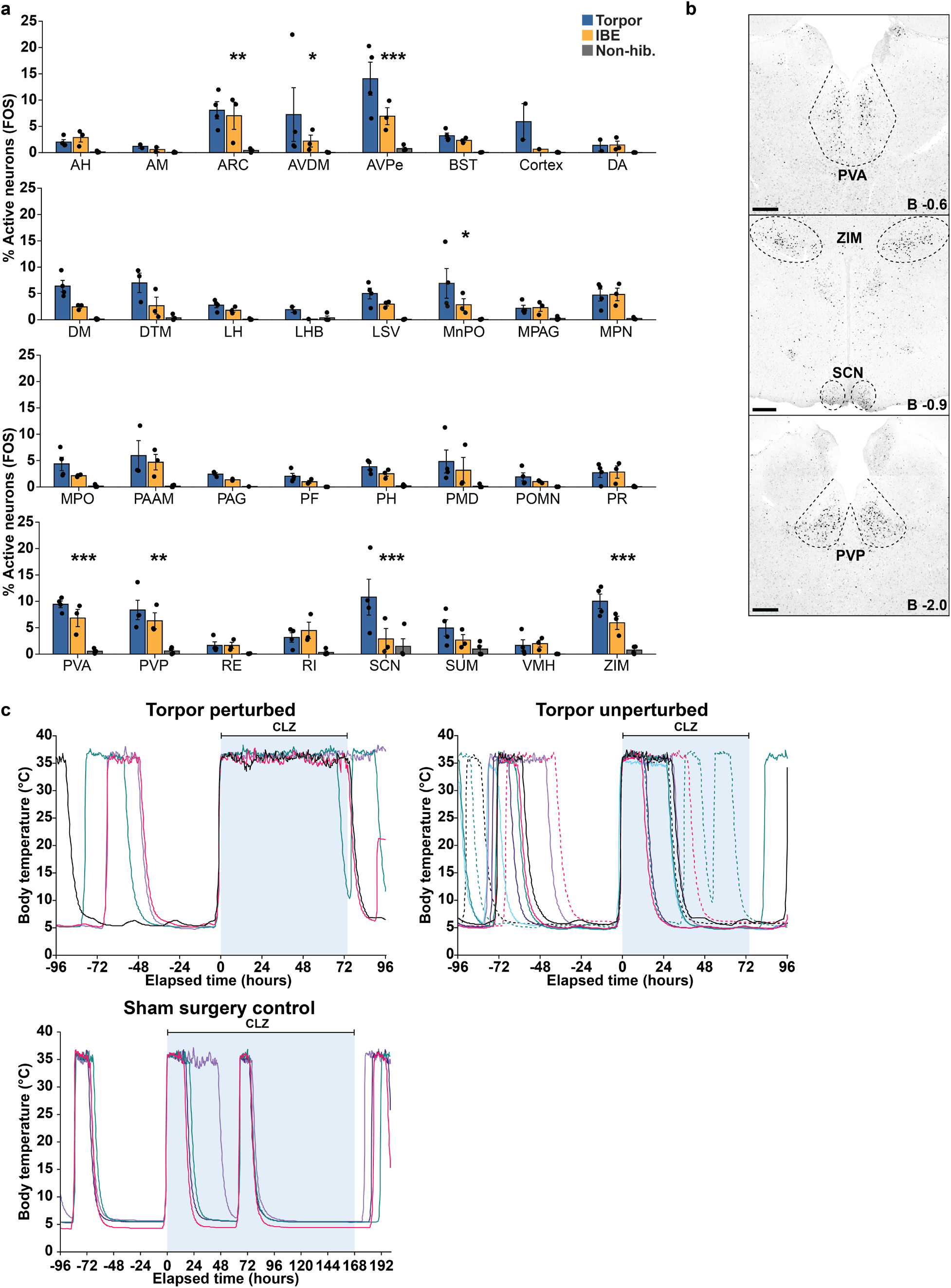
Hibernation-associated neuronal activity and disruption of torpor entry during targeted silencing. **a**, Bar plots showing the percentage of FOS-expressing neurons across all surveyed regions (n = 32) and hibernation stages (non-hibernating = 3 animals, IBE = 3 animals, torpor = 4 animals, p-adjusted < 0.05, **p-adjusted < 0.01, ***p-adjusted < 0.001). Data presented as mean ± s.e.m. **b**, Representative FOS immunostaining images of an animal in torpor. The top four deep torpor-active regions after the AVPe are shown: Paraventricular thalamic nucleus anterior (PVA), zona incerta (ZIM), suprachiasmatic nucleus (SCN), and paraventricular thalamic nucleus posterior (PVP). **c**, Hibernating hamsters (n = 13) were injected with AAV-hSyn-hM4D(Gi)-mCherry, targeting different areas of the anterior hypothalamus. After recovery of DT bouts, clozapine (CLZ, 5 mg/kg) was administered via drinking water. Hamster core body temperature traces are shown, demonstrating CLZ-dependent perturbation of torpor activity in a subset of animals (n = 4) while torpor entry remains unaffected in other animals (n = 9).

**Extended Data Fig. 3.**
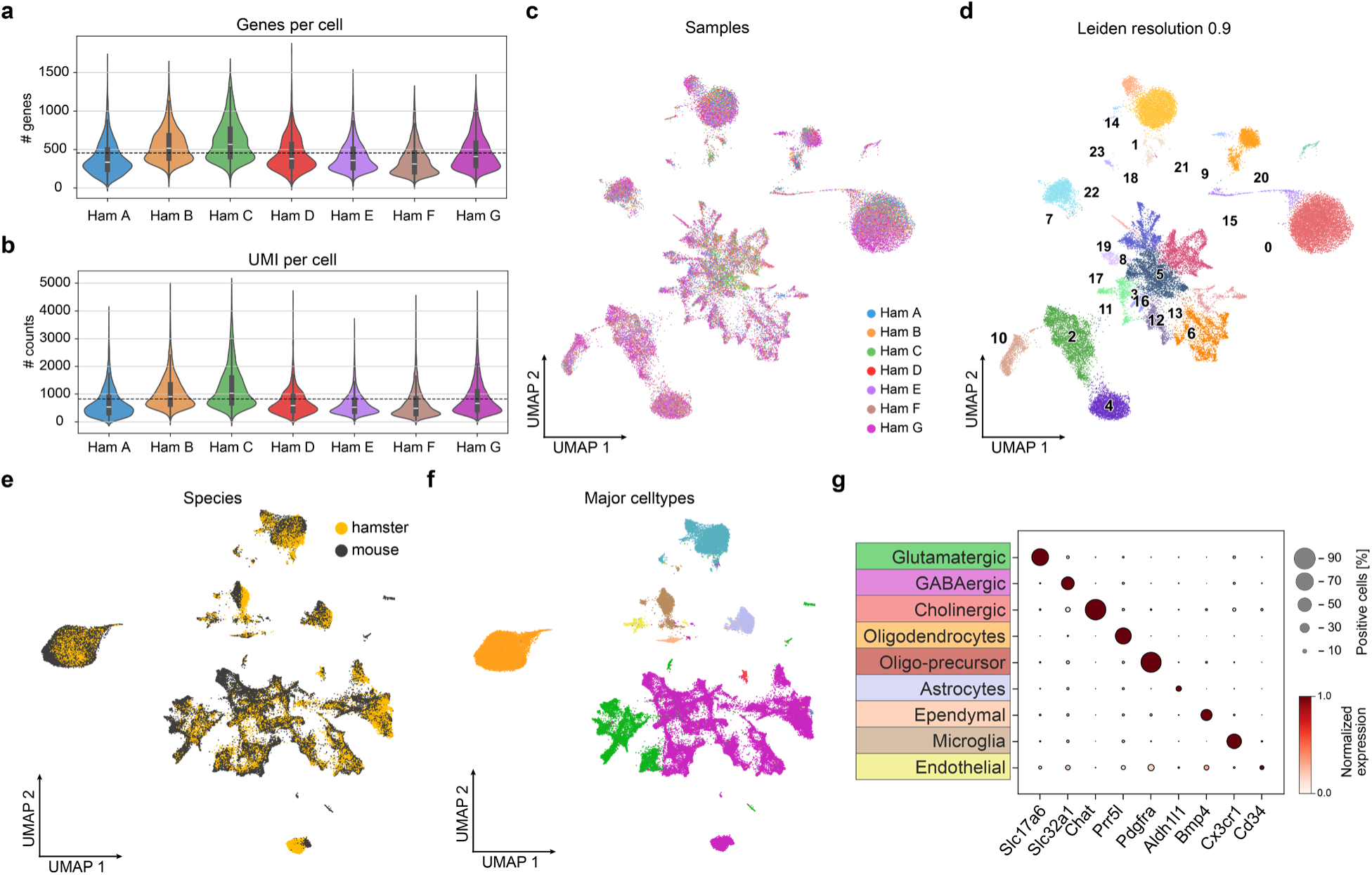
snRNA-seq analysis of hamster and mouse POA and separation of major neuronal and non-neuronal cell types. Our snRNA-seq and snATAC-seq data from hamster POA samples were first aligned with the mouse POA multiome dataset from Kaplan et. al^75^, separating major neuronal and non-neuronal cell types from both datasets. **a,b**, Violin plots showing the number of detected genes (**a**) and unique transcripts (UMIs) (**b**) per cell in each hamster snRNA-seq sample. **c,d**, UMAP plots showing 43,243 nuclei across 7 hamster samples and their Leiden clustering. **e,f**, UMAP plots showing an integrated dataset containing 45,553 hamster and mouse nuclei, and their separation into major cell types. **g**, Dot plot showing the expression of select gene markers of each major neuronal and non-neuronal cell type. Color indicates gene expression scaled across different cell types. Circle size indicates the percentage of cells expressing each marker gene.

**Extended Data Fig. 4.**
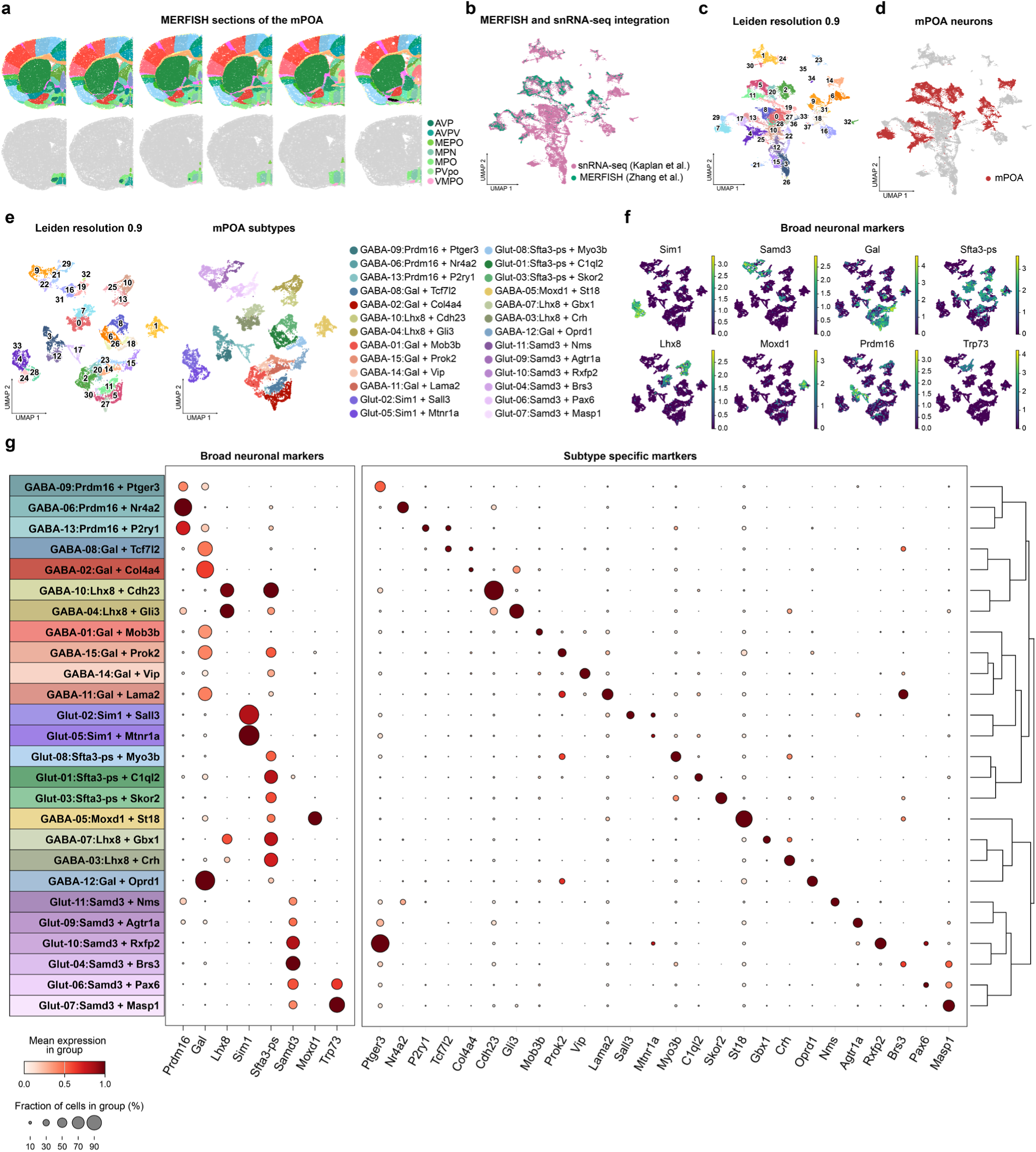
Creation of a mouse mPOA reference transcriptomic dataset. A mouse reference transcriptomic dataset restricted to cell types in our target regions – the medial and median POA (mPOA) – was generated. To this end, mouse spatial transcriptomic data (MERFISH)^76^ was subset to only include neurons found in the mPOA and co-clustered with mouse neuronal snRNA-seq populations from Extended Data Fig. 3f, g, enabling the annotation of 15 GABAergic and 11 glutamatergic mPOA neuronal populations in the mouse snRNA-seq dataset. **a**, Coronal brain sections indicating subregions of the mPOA. MERFISH data was subset to only include neurons within the indicated mPOA regions. **b**, UMAP plot showing the integrated MERFISH mPOA and mouse snRNA-seq dataset. **c,d**, UMAP plot showing Leiden clustering (**c**) and regional annotation (**d**) of the integrated MERFISH mPOA and mouse snRNA-seq dataset. **e**, UMAP plot of only mPOA neurons from (d) colored by Leiden cluster number (left), or annotated clusters (right). Annotation comprises cell class (Glut = glutamatergic, GABA = GABAergic), followed by key marker genes. These cells comprise the mouse mPOA reference transcriptomic dataset. **f**, UMAP plot showing expression of select marker genes in mouse mPOA. **g**, Dot plot showing the expression of cell-type-specific marker genes in mouse mPOA.

**Extended Data Fig. 5.**
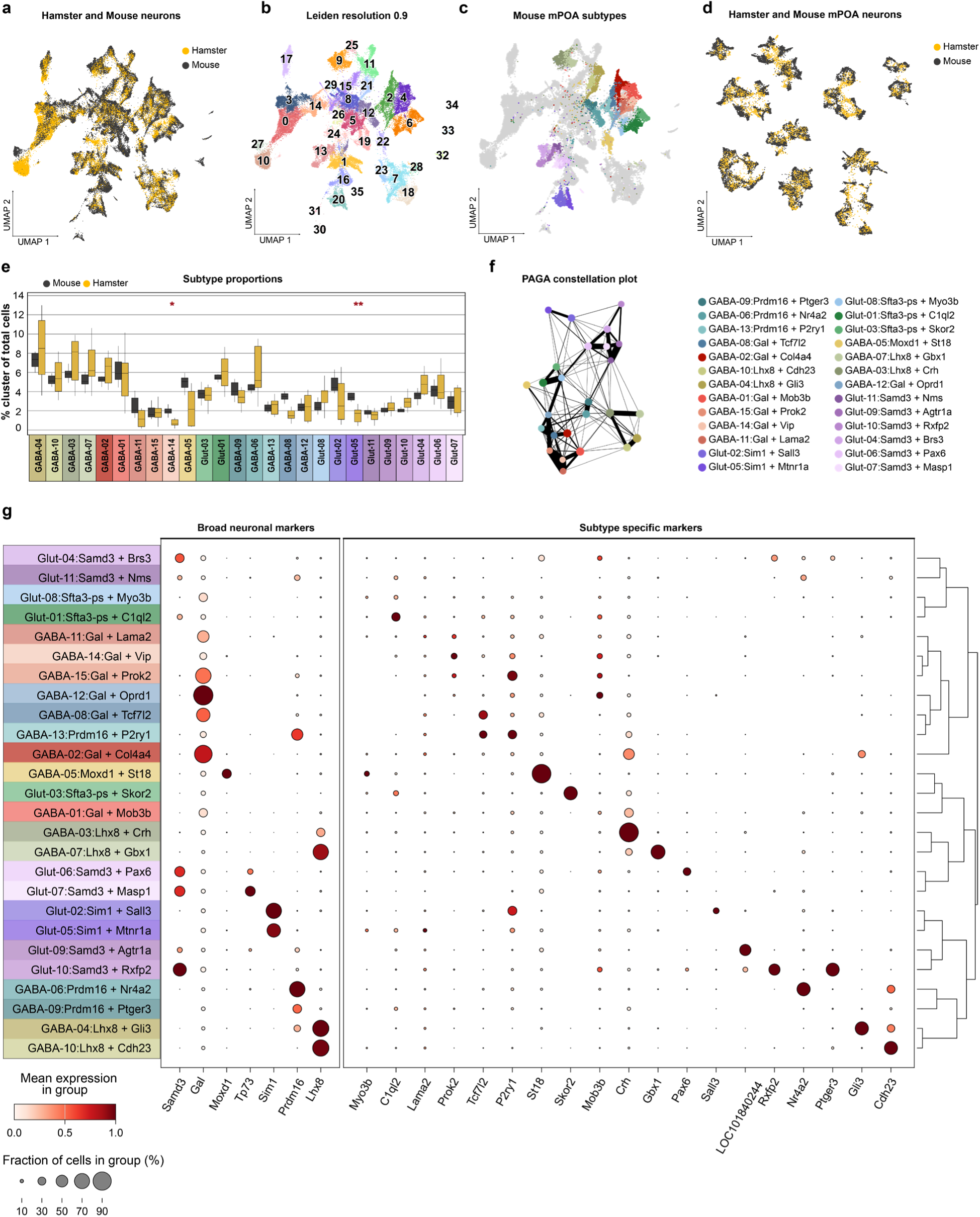
Integration of hamster snRNA-seq with the mouse mPOA reference transcriptomic dataset. Leveraging the annotations from the mouse mPOA reference transcriptomic dataset (Extended Data Fig. 4), we annotated neurons from the hamster POA snRNA-seq dataset to generate the final cross-species atlas. **a,b**, UMAP plot showing integrated hamster and mouse neuronal snRNA-seq data, colored by species of origin (**a**), or Leiden cluster (**b**). **c**, UMAP plot highlighting the location of mouse mPOA neurons, colored based on clusters from the mouse mPOA reference transcriptomic dataset. **d**, UMAP plot as in Fig. 3 displaying all mouse and hamster mPOA neurons from the snRNA-seq dataset, colored by the clusters defined in the mouse mPOA reference transcriptomic dataset. **e**, Relative abundance of each cell type in the mouse and hamster POA dataset. Three cell types show significant differences in abundance (* P < 0.05, ** P < 0.01, Mann-Whitney U test). **f**, Constellation plot showing the relationships between subtypes based on their latent space. **g**, Dot plot showing the expression of cell-type-specific marker genes for hamster neurons.

**Extended Data Fig. 6.**
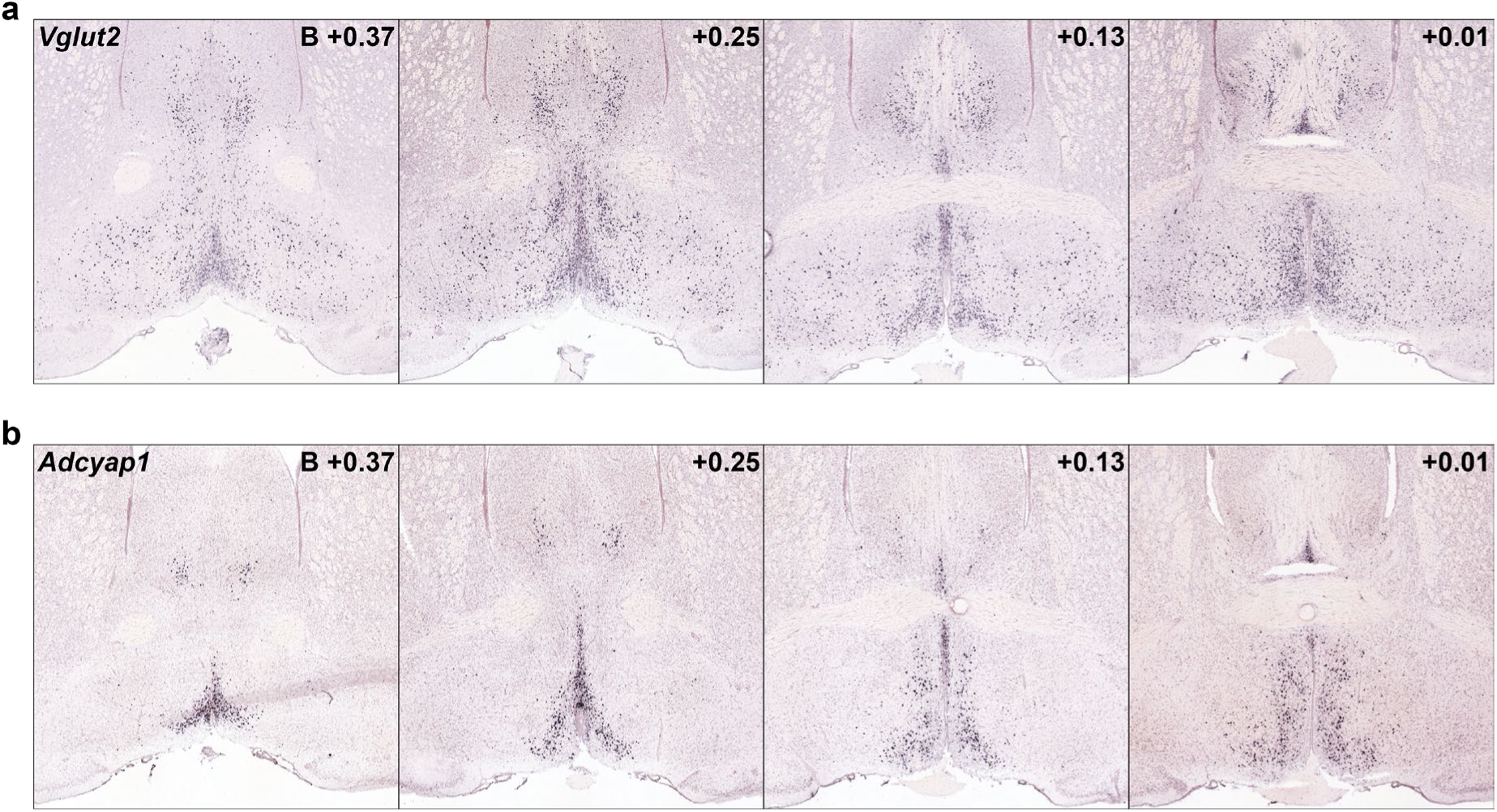
Anatomical distribution of mouse FIT-associated neurons. **a,b**, Example coronal sections of the mouse brain containing the anterior POA, showing ISH signal for marker genes *Slc17a6/Vglut2* (A) and *Adycap1* (B). Data from mouse.brain-map.org^103^.

**Extended Data Fig. 7.**
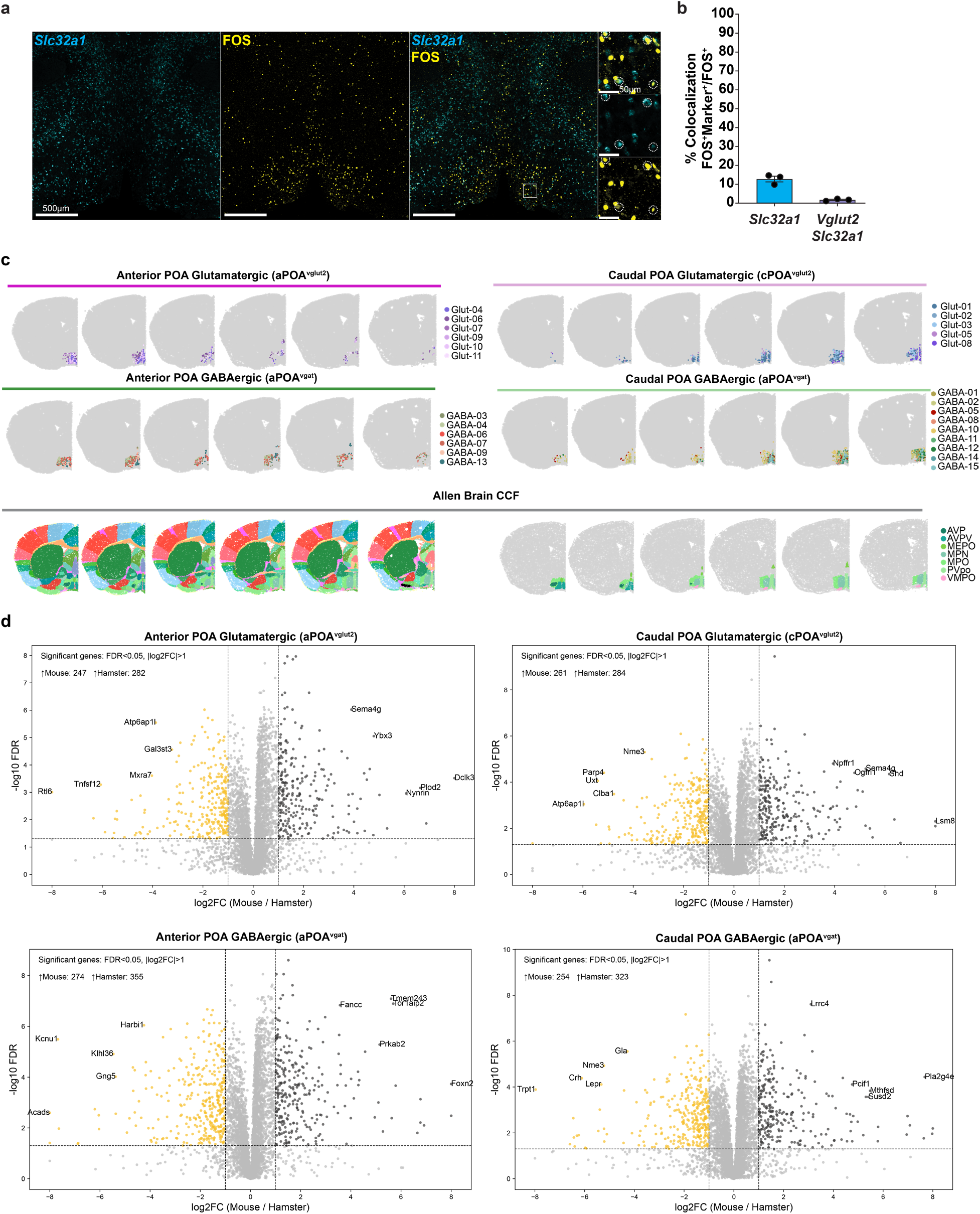
Marker expression of DT-active neurons; anatomical localization and cross-species differential gene expression of POA neuronal populations. **a**, Example coronal sections showing the aPOA of a Syrian hamster collected during DT onset. Immunofluorescent staining against FOS indicates the location of active neurons (yellow), whereas ISH indicates the expression of the marker gene *Slc32a1* (*Vgat*). High-magnification images are shown as insets with example FOS+ *Slc32a1*+ cells circled. **b**, Quantification of the fraction of FOS+ neurons during torpor that express *Slc32a1* (12.8% ± 1.5, animals = 3) and dual-expressing *Slc17a6/Slc32a1* (1.8% ± 0.3, n = 3) neurons. Data presented as mean ± s.e.m. **c**, MERFISH plot showing the spatial distribution of neuronal subtypes in the mouse POA separated by their major cell types and anatomical location. **d**, Differential gene expression between hamster and mouse neurons across the four major neuron groups.

**Extended Data Fig. 8.**
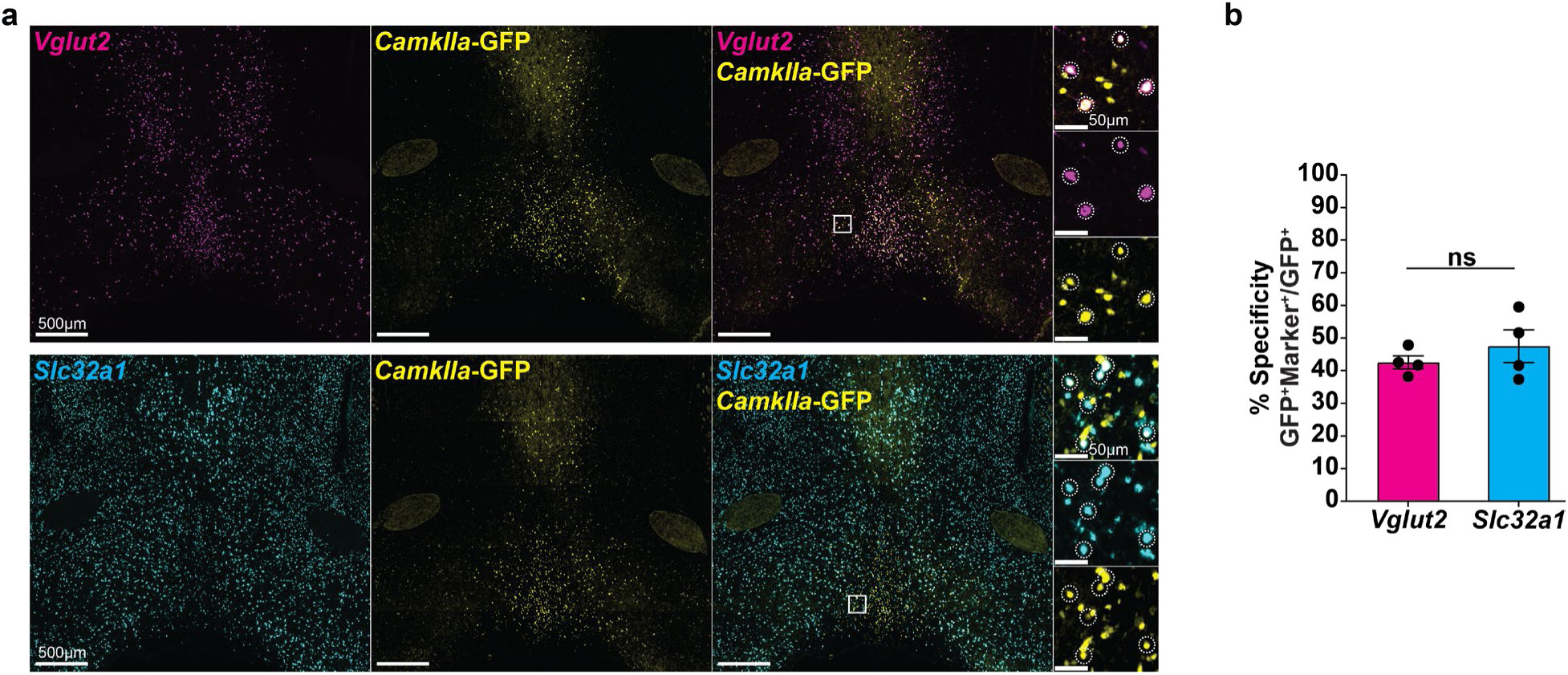
*CamkIIa* promoter-based AAVs lack specificity for glutamatergic neurons in the hamster aPOA. Hamsters (n = 4) were stereotaxically administered AAV8-CamkIIa-GFP and harvested ∼2 weeks later for immunostaining and ISH. **a**, Example coronal sections showing the hamster POA comprising AVPe, MPA, and MnPO. Immunofluorescent staining against GFP indicates the location of virally labeled cells (yellow), whereas ISH indicates the expression of genes *Slc17a6* (*Vglut2; magenta*) and *Slc32a1 (Vgat; cyan)*. High-magnification images are shown on the right with example GFP+ *Marker*+ cells circled. **b**, Quantification of the fraction of AAV-CamkIIa-GFP+ neurons that express *Slc17a6* (42.6% ± 2.0, animals = 4), and *Slc32a1* (47.5 ± 5.0%, animals = 4). Data presented as mean ± s.e.m.

**Extended Data Fig. 9.**
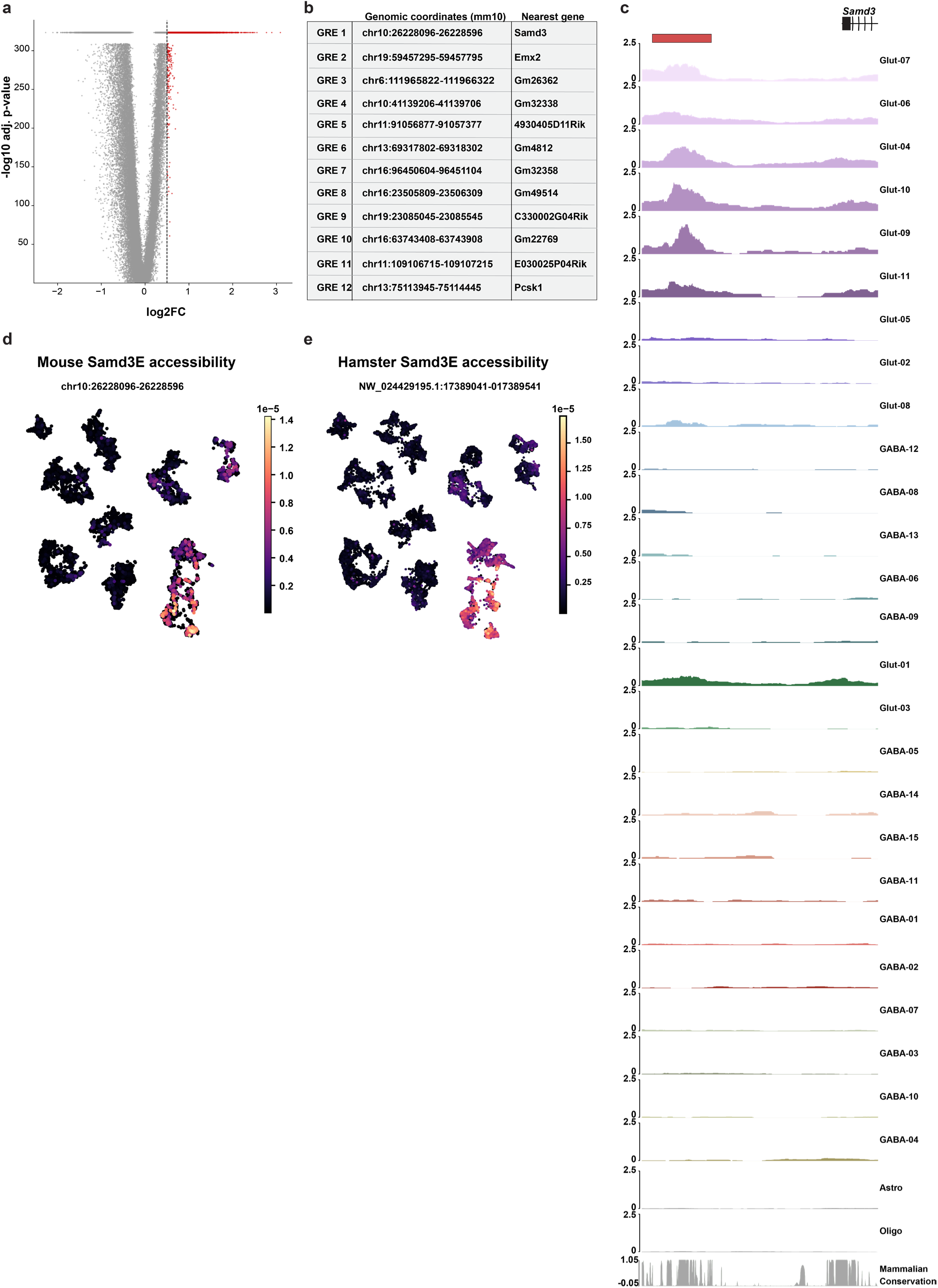
snATAC-seq analysis reveals *Samd3E* as tool to target POA neurons. **a**, Differential accessibility analyses of genomic elements in aPOA^Vglut2^ neurons vs. all other neurons in mice; top regions (log2FC > 0.5) are marked in red for further analysis. **b**, Genomic location and nearest gene of the top candidate putative GREs. **c**, ATAC-seq genome browser tracks (normalized counts per location) of individual mouse neuronal subtypes showing the mouse *Samd3*-proximal putative GRE (*Samd3E,* red box), with accompanying sequence conservation below. **d,e**, UMAP feature plots depicting *Samd3E* accessibility in the indicated species.

**Extended Data Fig. 10.**
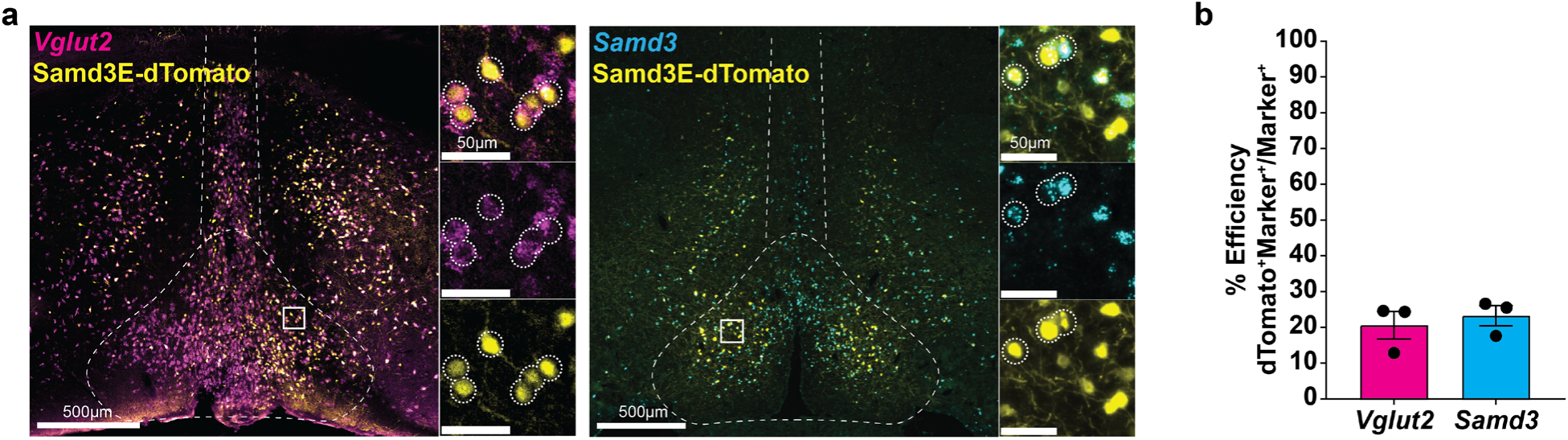
Samd3E-AAV transduction efficiency. **a**, Efficiency of targeting aPOA^Vglut2^ and aPOA^Samd3^ neurons with AAV8-Samd3E-dTomato. Syrian hamsters (n = 3) were injected with AAV8-Samd3E-dTomato. Immunofluorescent staining against dTomato was used to identify virally labeled cells (yellow), whereas ISH indicates the expression of the marker gene *Slc17a6* (*Vglut2*, magenta) and *Samd3* (cyan), as in Fig. 4g. **b**, Quantification of the fraction of *Slc17a6*+ cells that also express dTomato+ (20.6 ± 3.9%, animals = 3) and *Samd3*+ cells that also express dTomato+ (23.3 ± 2.8%, animals = 3). Given limited viral spread, this quantification was limited to the area of viral expression within the aPOA. These results indicate that AAV8-Samd3E-dTomato achieves high specificity and modest efficiency for targeting aPOA^Vglut2^ and aPOA^Samd3^ neurons.

**Extended Data Fig. 11.**
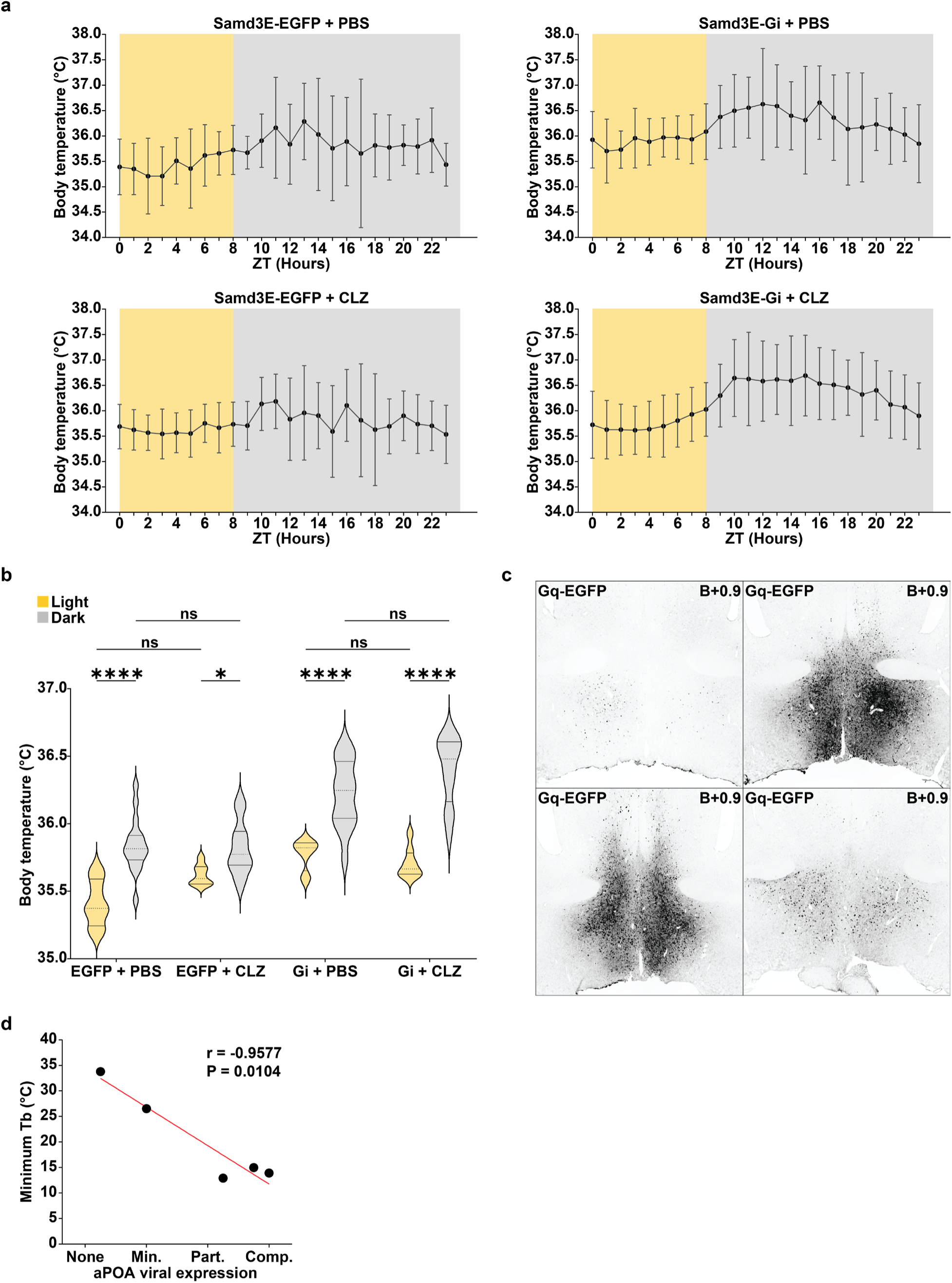
Circadian T_b_ patterns and correlation between viral aPOA expression and T_b_ decrease. **a**, Circadian fluctuation of T_b_ in Samd3E-EGFP- and Samd3E-Gi-DREADD-injected hamsters following administration of CLZ (5 mg/kg) or control PBS. Light and dark cycle phases are represented by yellow and grey, respectively. Data presented as mean ± s.d. of core body temperatures during each hour. **b**, Violin plot quantification of body temperature range between light (yellow) and dark (grey) phases of Samd3E-EGFP-(n = 5 animals) or Samd3E-Gi-DREADD-injected animals (n = 12) following CLZ or control PBS administration (Tukey’s multiple comparisons test; *P-adjusted < 0.05, ****P-adjusted < 0.0001, ns = not significant P > 0.05). **c**, Immunofluorescence images in the aPOA of additional Samd3E-Gq-DREADD-EGFP-injected animals. **d**, Correlation between minimum T_b_ reached following CLZ administration and viral expression within the aPOA (n = 5 animals, r = -0.95, P = 0.01).

**Extended Data Fig. 12.**
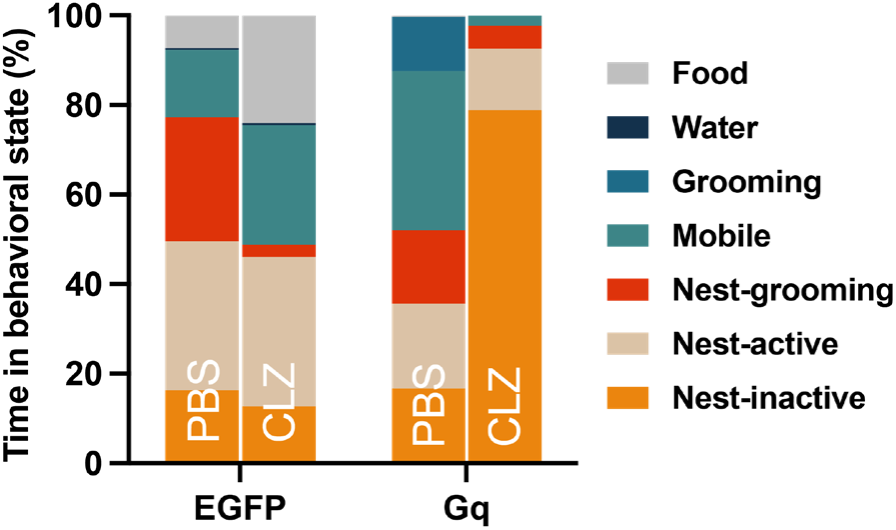
Changes in behavioral repertoire after aPOA^Vglut2^ neuronal activation. Percentage of time in behavioral states over a 10 min observation window post-injection with PBS or CLZ across all hamsters injected with AAV-Samd3E-EGFP-P2A-Synaptophysin-mRuby (n = 3) and AAV-Samd3E-hM3D(Gq)-nlsGFP (n=6).

**Extended Data Fig. 13.**
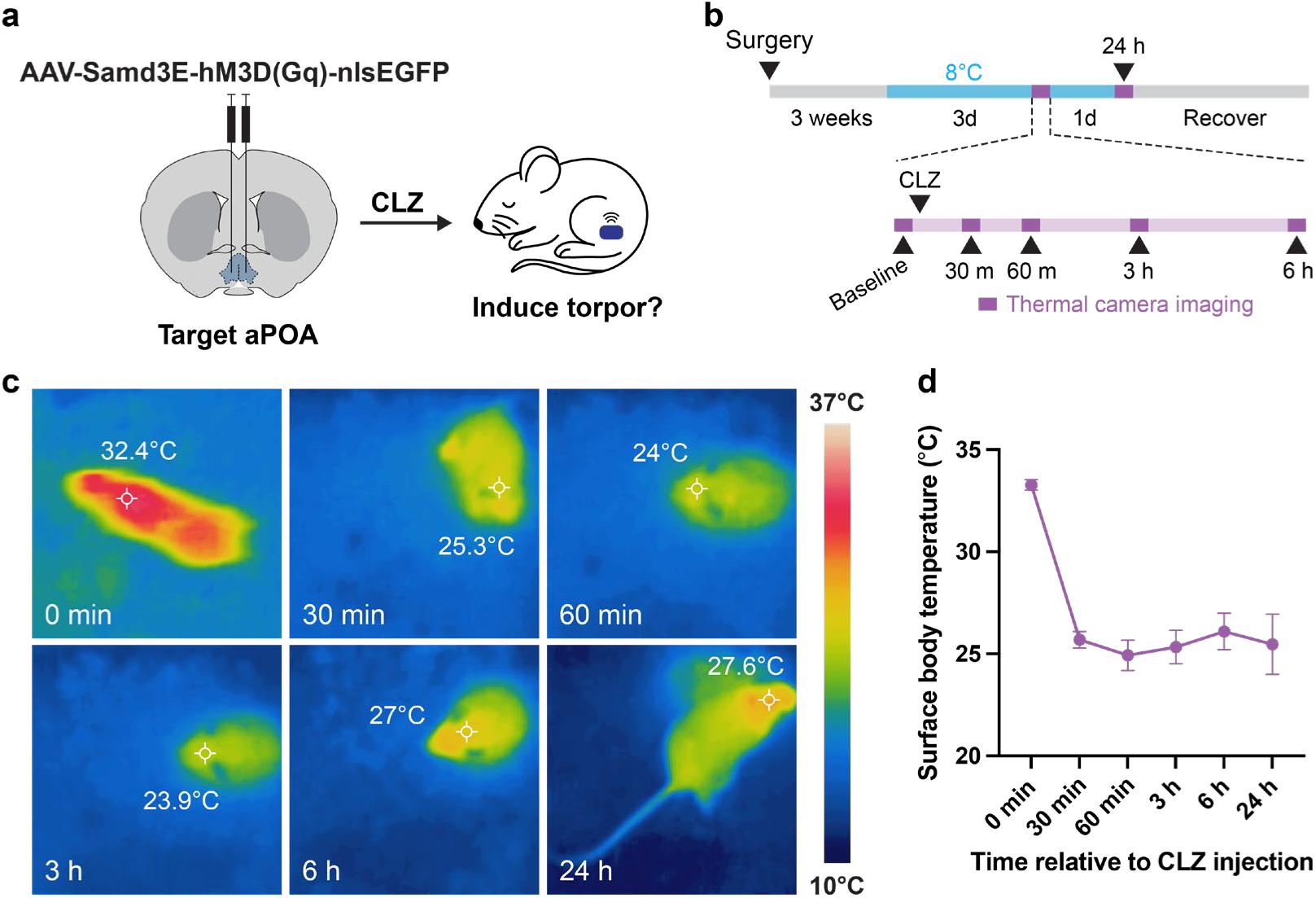
Stimulation of aPOA^Vglut2^ neurons in mice and resulting body temperature change. **a**, Schematic of experimental design. **b**, Timeline of experiment and analysis. Mice were acclimated to 8°C for 3 days before injection with CLZ (5 mg/kg) and imaged with a thermal camera at baseline, 30 min, 60 min, 3 hours, 6 hours, and 24 hours post injection at 8°C. **c**, Representative thermographic images obtained by chemogenetic stimulation of aPOA^Vglut2^ neurons. **d**, Surface body temperature before (0 min) and after (30 min to 24 h) CLZ injection (n=7 mice). Data are presented as mean ± s.e.m.

**Extended Data Fig. 14.**
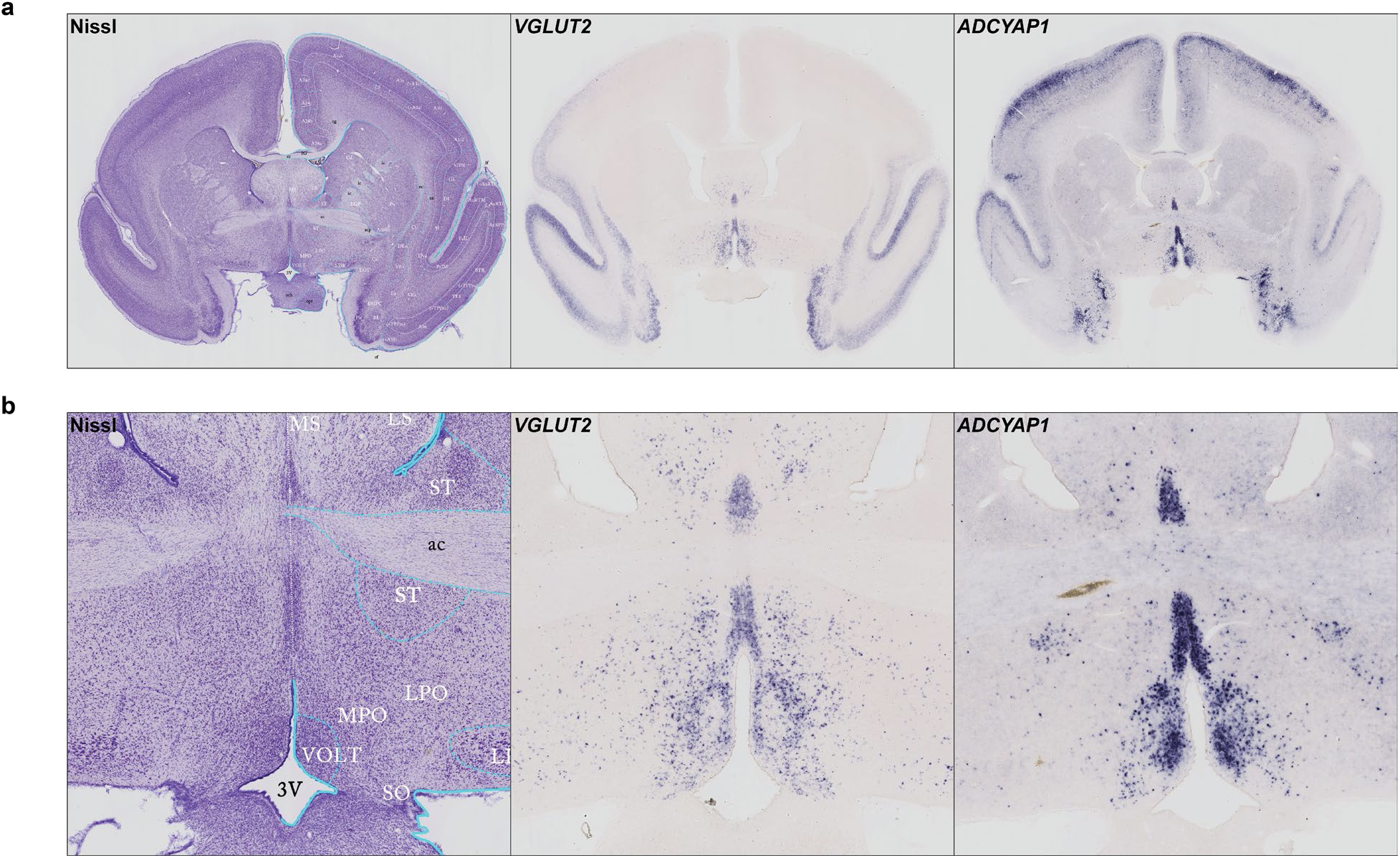
*In situ* hybridization for marker of torpor-associated genes in marmosets. **a**, Example coronal sections of the marmoset brain containing the anterior POA showing ISH signal for marker genes *Slc17a6/Vglut2* and *Adycap1*. **b**, Higher resolution images from (A) displaying the anterior POA. Data from https://gene-atlas.brainminds.jp/^90–92^.

